# Structural basis of long-range transcription-translation coupling

**DOI:** 10.1101/2024.07.20.604413

**Authors:** Chengyuan Wang, Vadim Molodtsov, Shashank Shandilya, Linlin You, Jing Zhang, Konstantin Kuznedelov, Bryce E. Nickels, Jason T. Kaelber, Gregor Blaha, Richard H. Ebright

**Author notes:** Contributed equally (Chengyuan Wang, Vadim Molodtsov).

## Abstract

Structures recently have been reported of molecular assemblies that mediate transcription-translation coupling in *Escherichia coli*. In these molecular assemblies, termed “coupled transcription-translation complexes” or “TTC-B”, RNA polymerase (RNAP) directly interacts with the ribosome, the transcription elongation factor NusG or its paralog RfaH forms a bridge between RNAP and ribosome, and the transcription elongation factor NusA optionally forms a second bridge between RNAP and ribosome. Here, we report structures of coupled transcription-translation complexes having mRNA spacers between RNAP and ribosome longer than the maximum-length mRNA spacer compatible with formation of TTC-B. The results define a new class of coupled transcription-translation complex, termed “TTC-LC,” where “LC” denotes “long-range coupling.” TTC-LC differs from TTC-B by a ∼60° rotation and ∼70 Å translation of RNAP relative to ribosome, resulting in loss of direct interactions between RNAP and ribosome and creation of a ∼70 Å gap between RNAP and ribosome. TTC-LC accommodates long mRNA spacers by looping out mRNA from the gap between RNAP and ribosome. We present evidence that TTC-LC is an intermediate in assembling and disassembling TTC-B, mediating pre-TTC-B transcription-translation coupling before a ribosome catches up to RNAP, and mediating post-TTC-B transcription-translation coupling after a ribosome stops moving and RNAP continues moving. We show that TTC-B, but not TTC-LC, is severely defective in intrinsic, RNA-hairpin-dependent termination, and that both TTC-B and TTC-LC are severely defective in Rho-dependent termination.

## Introduction

Bacterial transcription and translation occur in the same cellular compartment^1–2^, occur at the same time^1–2^, and, in many cases, are functionally coordinated and physically coupled processes, in which the rate of transcription by the first molecule of RNA polymerase (RNAP) synthesizing an mRNA is coordinated with the rate of translation by the first ribosome translating the mRNA^2–14^. Transcription-translation coupling increases transcription elongation rates, suppresses transcription pausing, suppresses transcription termination, increases translation elongation rates, suppresses translation pausing, and plays roles in regulation of gene expression^3–24^. In the bacterium *Escherichia coli*, the physical coupling of transcription and translation is mediated by the transcription elongation factor NusG and its paralog RfaH^4–14, 25–29^. NusG is a general coupling factor that couples transcription and translation at most genes in *E. coli*^4–14, 25–29^, and RfaH is a specialized, regulon-specific coupling factor that couples transcription and translation at a small number of genes in *E. coli* that contain a DNA sequence, termed an *ops* site, required for loading RfaH onto RNAP^5–7, 10–13, 25–29^. NusG and RfaH each consists of an N-terminal domain (NusG-N or RfaH-N) that interacts with RNAP, a C-terminal domain (NusG-C or RfaH-C) that interacts with ribosomal protein S10, and an interdomain linker^4–5, 7–14, 25–29^. *E. coli* transcription-translation coupling is further mediated by a second transcription elongation factor, NusA^30^.

Structures of NusG- and RfaH-containing *E. coli* transcription-translation complexes (TTCs) recently have been reported^31–35^. Two distinct structural and functional classes of TTCs have been described: (i) “collided TTCs”, also referred to as “TTC-A”, which have structural features that suggest they are translationally inactive, or only partly, active products of collision between a ribosome and RNAP^31–35^; and (ii) “coupled TTCs”, also referred to as “TTC-B,” which have structural features that suggest that they are the translationally active complexes that mediate functional transcription-translation coupling^32–35^. Structures of NusG- and RfaH-coupled TTC-B are superimposable, differing only in that NusG-coupled TTC-B shows greater conformational heterogeneity in the orientation of RNAP relative to the ribosome than RfaH-coupled TTCs (Fig. 1)^34^. In both NusG- and RfaH-coupled TTC-B, the coupling factor NusG or RfaH bridges RNAP and the ribosome, with NusG-N or RfaH-N interacting with RNAP, and with NusG-C or RfaH-C simultaneously interacting with ribosomal protein S10 in the ribosome 30S subunit^32–35^. When present, the coupling factor NusA forms a second bridge between RNAP and the ribosome; this second bridge supplements, but cannot substitute for, the bridge formed by NusG or RfaH (Fig. 1)^33–35^.

**Fig. 1.**
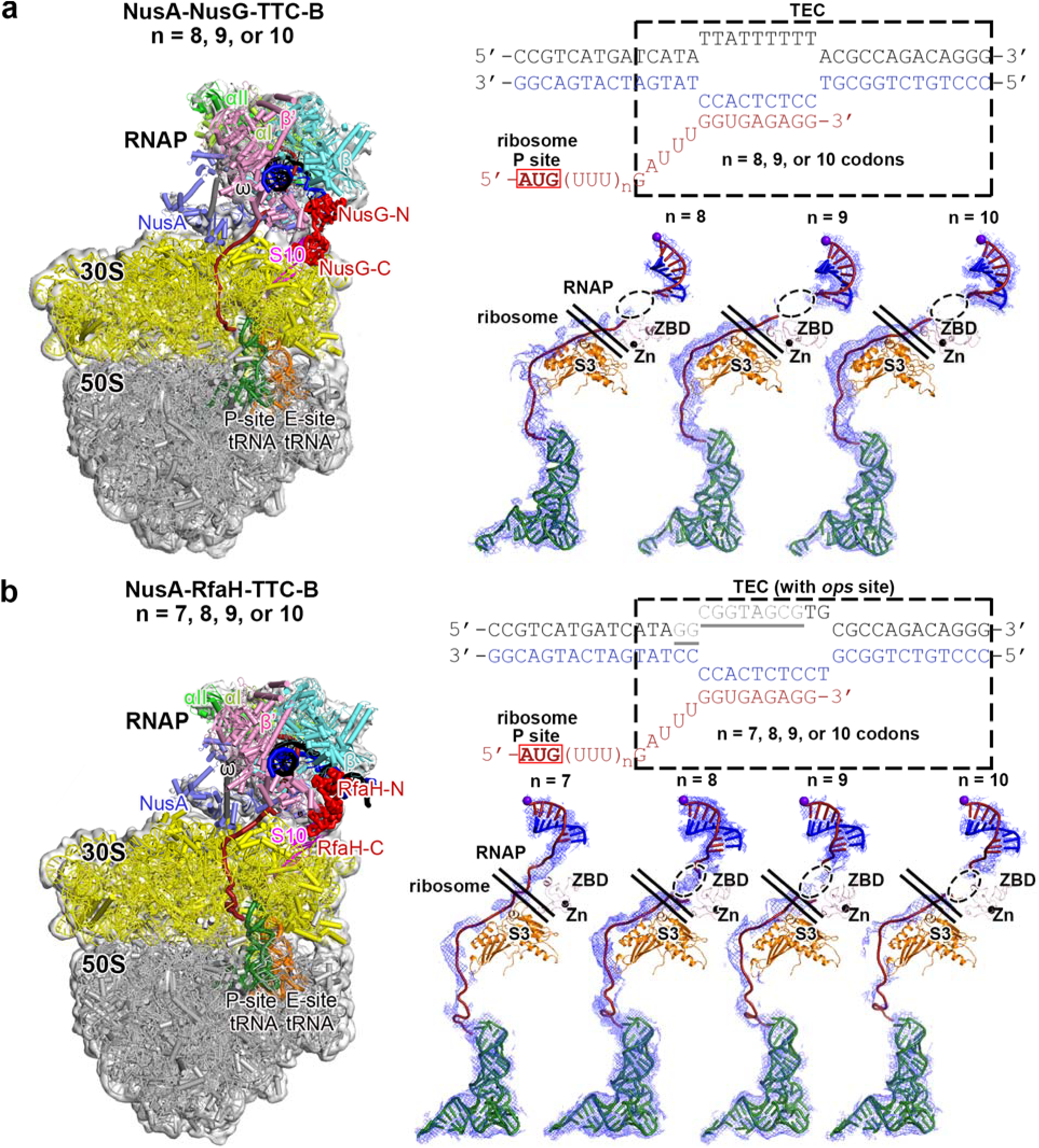
Structures of NusG-, NusA-coupled TTC-B and RfaH-, NusA-coupled TTC-B. **(a)** NusA-NusG-TTC-B (PDB 6X6T)^33^. Left: structure. Top right: nucleic-acid scaffold used for structure determination. Bottom right: accommodation of different mRNA spacer lengths in NusA-NusG-TTC-B. **(b)** NusA-RfaH-TTC-B (PDB 8UQP).^34^ Left: structure. Top right: nucleic-acid scaffold used for structure determination (*ops* site in grey and underlined). Bottom right: accommodation of different mRNA spacer lengths in NusA-RfaH-TTC-B. Left panels show EM density (grey surface) and fit (ribbons) for TEC, NusG or RfaH, and NusA (at top), and for ribosome 30S and 50S subunits and P-site and E-site tRNAs (at bottom). RNAP β’, β, α^1^, α^2^, and ω subunits are in pink, cyan, light green, green, and grey; 30S subunt, 50S subunit, P-site tRNA, and E-site tRNA are in yellow, grey, green, and orange; DNA non-template strand, DNA template strand, and mRNA are in black, blue, and brick-red; NusG or RfaH, NusA, and ribosomal protein S10 are in red, light blue, and magenta; ribosome L7/L12 stalk is omitted for clarity in this and subsequent images. Top right subpanels show nucleic-acid scaffolds used for structure determination, Each scaffold comprises non-template-strand oligodeoxyribonucleotide (black), template-strand oligodeoxyribonucleotide (blue), and mRNA oligoribonucleotide (brick red). Dashed box labeled “TEC,” portion of nucleic-acid scaffold that forms TEC upon addition of RNAP (10 nt non-template- and template-strand ssDNA segments forming “transcription bubble,” 9 nt of mRNA engaged with templat-strand DNA as RNA-DNA “hybrid,” and 5 nt of mRNA, on diagonal, in RNAP RNA-exit channel); red box labeled “ribosome P-site,” mRNA AUG codon intended to occupy ribosome active-center P-site upon addition of ribosome and tRNA^fMet^. Bottom right subpanels show accommodation of different mRNA spacer lengths inside complex. EM density, blue mesh; mRNA, brick red (disordered mRNA nucleotides indicated by dashed ovals); template-strand DNA in RNA-DNA hybrid, blue; RNAP active-center catalytic Mg^2+^, purple sphere; ribosomal protein S3 in ribosome 30S subunit, orange; tRNA in ribosome P-site, green. Pairs of diagonal black lines indicate edges of RNAP and ribosome.

Structures of NusG- and RfaH-coupled TTC-B have been obtained using nucleic-acid scaffolds for mRNA spacers between RNAP and ribosome that were 7, 8, 9, or 10 codons in length (21, 24, 27, or 30 nt in length) (Fig. 1)^32–35^. Structures of TTC-B having mRNA spacers 7, 8, 9, or 10 codons in length are superimposable^32–35^, with the differences in mRNA spacer length being accommodated by different extents of compaction and disorder of mRNA in the RNAP-exit channel (Fig. 1)^32–35^. Based on the volume of the RNAP RNA-exit channel, the RNAP-exit channel is predicted to be able to accommodate a maximum of 5 codons (15 nt) of compacted and disordered mRNA ^33^; therefore, it is expected that TTC-B is compatible with, and can be formed, only for mRNA spacers of lengths ranging from 7 codons (21 nt) to 7 + 5 = 12 codons (36 nt)^33^.

Qureshi and Duss^36^ recently reported single-molecule fluorescence-resonance energy-transfer studies of *E. coli* transcription-translation coupling. Their results confirm the existence of functional and physical coupling between RNAP and ribosome, confirm the role of coupling factors NusG and NusA in enhancing coupling, and indicate, surprisingly, that coupling can occur not only over the range of RNAP-ribosome spacings that can be accommodated in TTC-B, but also over much greater RNAP-ribosome spacings. Based on their findings, Qureshi and Duss conclude that there exists a structural class of TTCs that can accommodate very long mRNA spacers between RNAP and ribosome, accommodating these spacers by looping out of the mRNA spacer between RNAP and ribosome. Qureshi and Duss further conclude that the long-range coupling, although it clearly occurs, is less stable than short-range coupling. The lower stability is reflected in a smaller number of complexes observed with long-range coupling than with short-range coupling.

Here, in order to define the structural basis of transcription-translation coupling with long mRNA spacers between RNAP and ribosome, and to test the proposal of looping out of long mRNA spacers between RNAP and ribosome, we determined cryo-EM structures of *E. coli* NusG- and RfaH-coupled TTCs with mRNA spacers longer than those that can be accommodated within TTC-B.

### Structure determination

We assembled TTCs using synthetic nucleic-acid scaffolds analogous to those used previously to determine structures of NusG- and RfaH-coupled TTCs (Figs. 1, 2b, 3b)^32–35^. The scaffolds consist of three parts: (i) DNA and mRNA determinants that direct formation of a transcription elongation complex (TEC) upon addition of RNAP; (ii) an mRNA AUG codon that directs formation of a translation complex having the AUG codon in the ribosome active-center product site (P-site) upon addition of a ribosome and tRNA^fMet^; and (iii) an mRNA spacer having, in intial studies, a length, *n*, of 12 or 13 codons, and in subsequent studies a length, *n*, of 17 or 20 codons (Figs. 2b, 3b). For determination of structures of RfaH-coupled TTCs, the nucleic-acid scaffolds also contained an *ops* site with consensus sequence and positioning relative to the unwound transcription bubble of the TEC (Fig 1b, right subpanel). For each nucleic-acid scaffold, we equilibrated the scaffold with RNAP to form a TEC; we than added coupling factor NusG or RfaH, and optionally also coupling factor NusA, to form a NusG- or RfaH-containing TEC; and we than added ribosomes and tRNA^fMet^ to form a NusG- or RfaH-containing TTC. For each resulting complex we then determined the structure using single-particle reconstruction cryo-EM (Extended Data Figs. 1-5; Extended Data Tables 1-2).

**Fig. 2.**
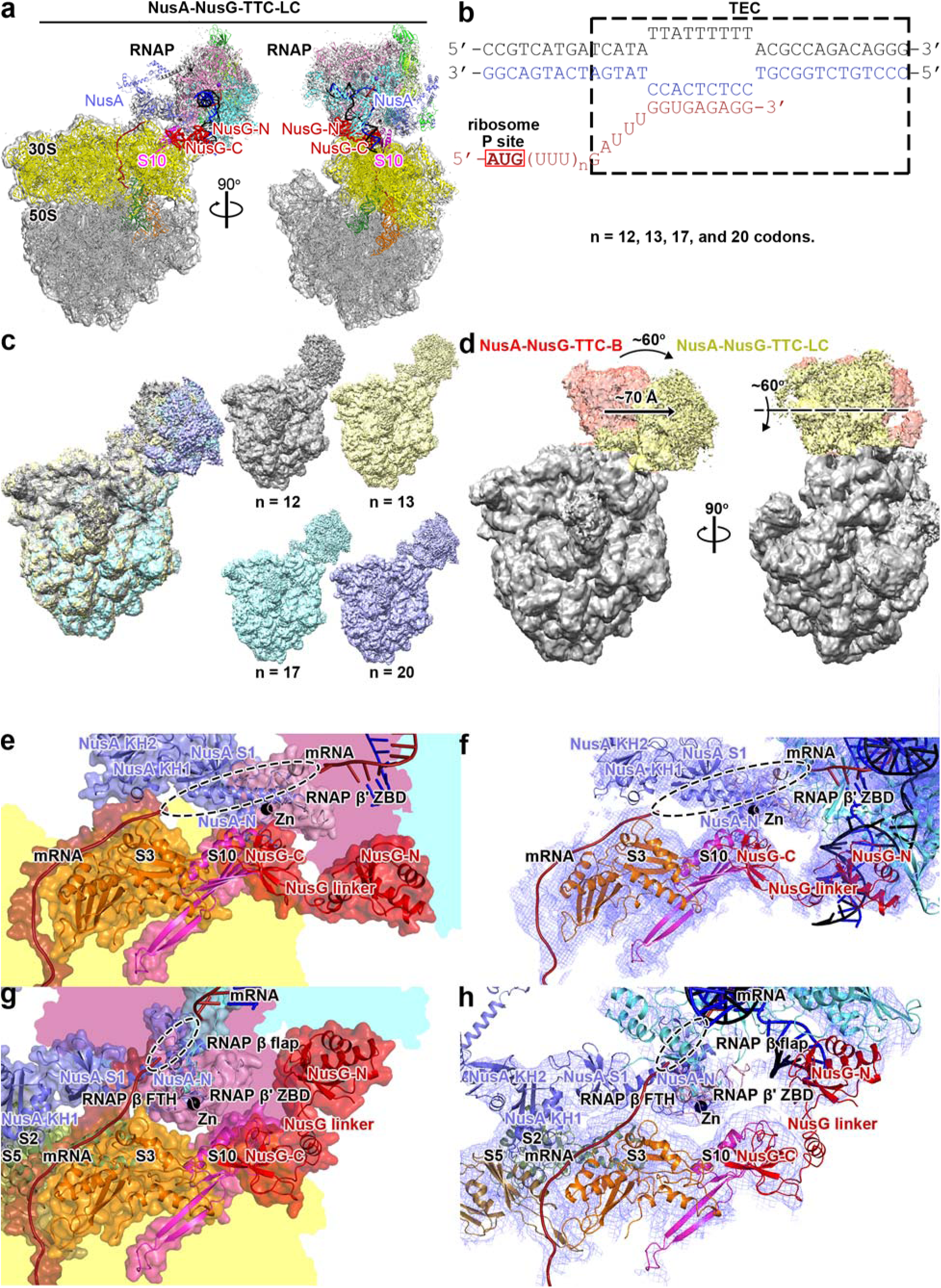
Structure of NusG-, NusA-coupled TTC-LC. **(a)** Structure of NusG-NusA-TTC-LC (n = 17, Extended Data Table 1). Two orthogonal views. Colors and features as in Fig. 1. **(b)** Nucleic-acid scaffold used for structure determination. Colors as in Fig. 1. **(c)** EM density maps for structures of NusG-NusA-TTC-LC obtained using nucleic-acid scaffolds with n = 12, 13, 17, and 20 (superimposition at left; individual EM maps and color scheme at right). View orientation as in (a). **(d)** Superimposition of NusA-NusG-TTC-LC (TEC in yellow and ribosome in grey) on NusA-NusG-TTC-B^33^ (subclass B1; TEC in pink and ribosome in grey). **(e)** RNAP-ribosome interface in NusA-NusG-TTC-LC (n = 17; identical interfaces for n = 12, 13, and 20). RNAP β’ zinc binding domain (ZBD), pink with Zn^2+^ ion as black sphere; NusG, red; NusA, slate blue; ribosomal protein S3, orange; ribosomal protein S10, magenta; DNA template strand, blue; mRNA, brick red; disordered mRNA segment, black dashed oval; portions of RNAP β’, β, and ribosome 30S not involved in the interactions are shaded pink, cyan, and yellow, respectively. In TTC-LC, NusG directly bridges RNAP and ribosome, with NusG-N interacting with RNAP, and NusG-C interacting with ribosomal protein S10. and NusA indirectly bridges RNAP and the mRNA segment interacting with ribosomal protein S3. **(f)** As (e), showing cryo-EM density as blue mesh. **(g)** RNAP-ribosome interface in NusA-NusG-TTC-B (n = 12)^33^. View orientation as in (e-f). In TTC-B, RNAP ZBD interacts with ribosomal protein S3; NusG directly bridges RNAP and ribosome, interacting with RNAP and ribosomal protein S2 (forest green) and S5 (brown); and NusA directly bridges RNAP and ribosome, interacting with RNAP and ribosomal protein S3. **(h)** As (f), but for NusA-NusG-TTC-B (n = 12)^33^.

**Fig. 3.**
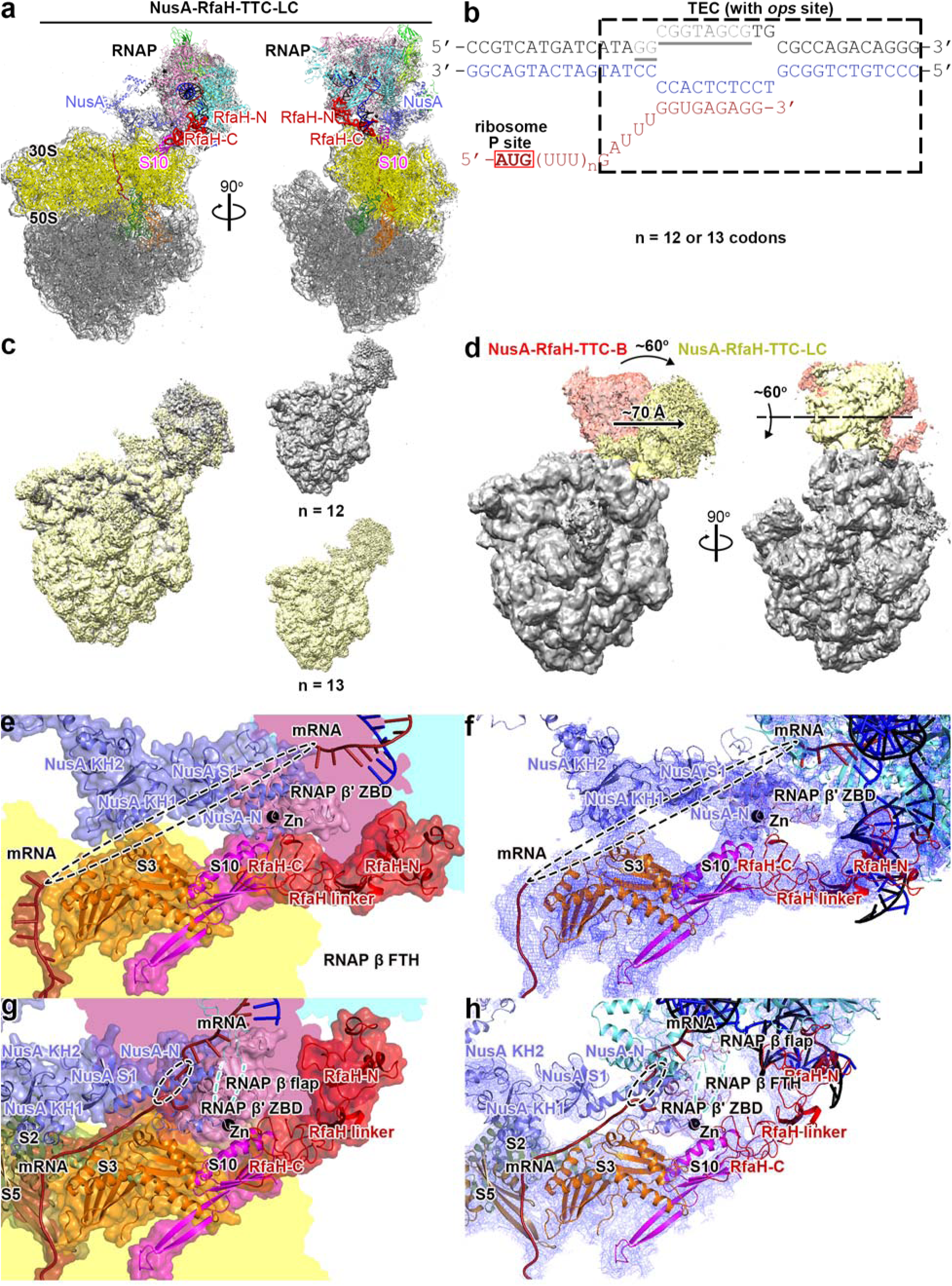
Structure of RfaH-, NusA-coupled TTC-LC. **(a)** Structure of RfaH-NusA-TTC-LC (n = 13, Extended Data Table 2). Views and colors as in Figs. 1 and 2a. **(b)** Nucleic-acid scaffold used for structure determination. Colors and features as in Fig. 1. **(c)** EM density maps for structures of RfaH-NusA-TTC-LC obtained using nucleic-acid scaffolds with n = 12 and 13 (superimposition at left; individual EM maps and color scheme at right). View orientation as in (a). **(d)** Superimposition of RfaH-NusA-TTC-LC (TEC in yellow and ribosome in grey) on RfaH-NusA-TTC-B^34^ (TEC in pink and ribosome in grey). **(e)** RNAP-ribosome interface in RfaH-NusA-TTC-LC (n = 13; identical interfaces for n = 12). RfaH, red. Other colors as in Fig. 2e. In TTC-LC, RfaH directly bridges RNAP and ribosome, with RfaH-N interacting with RNAP and RfaH-C interacting with ribosomal protein S10; and NusA indirectly bridges RNAP and ribosome, interacting with the mRNA segment that interacts with ribosomal protein S3. **(f)** As (e), showing cryo-EM density as blue mesh. **(g)** RNAP-ribosome interface in RfaH-NusA-TTC-B (n = 8)^34^. View orientation as in (e-f). In TTC-B, RNAP ZBD interacts with ribosomal protein S3; RfaH directly bridges RNAP and ribosome, interacting with RNAP and ribosomal protein S2 (forest green) and S5 (brown); and NusA directly bridges RNAP and ribosome, interacting with RNAP and ribosomal protein S3. **(h)** As (f), but for RfaH-NusA-TTC-B.

### TTC-LC: structures of NusG- and RfaH-coupled TTCs containing long--12 or 13 codons--mRNA spacers between RNAP and ribosome

We first determined structures of NusG- and RfaH-containing TTCs using nucleic-acid scaffolds having mRNA spacers of length, *n*, of 12 and 13 codons, representing mRNA spacers that are at or above the expected maximum length mRNA spacer that can be accommodated in, and can enable formation of, TTC-B (Figs. 2–3; Extended Data Figs. 1-3; Extended Data Tables 1-2). Using these scaffolds, we obtained structures of NusG- or RfaH-coupled, NusA-co-coupled TTCs that define a new structural class of TTCs, different from TTC-B (Figs 2-3). In this new structural class of TTCs, the position of RNAP relative to the ribosome differs from that in a NusG-coupled TTC-B subclass B1 and in RfaH-coupled TTC-B by a rotation of ∼60° and a translation of ∼70Å (Figs. 2d, 3d; Extended Data Figs. 1f, 2f, 3f) and differs from the orientation in NusG-coupled TTC-B subclasses B2 and B3 by a rotation of ∼45° and translation of ∼63Å and a rotation of ∼ 30° and translation of ∼47Å, respectively (Extended Data Fig. 1f). In this new structural class of TTCs, in contrast to in previously defined structural class TTC-B, there is no, or almost no, interaction between RNAP and the ribosome (Figs. 2e-h, 3e-h). The new structural class also differs from TTC-B in the path of mRNA from RNAP to ribosome, having a ∼70Å gap between mRNA in the RNAP RNA-exit channel and mRNA interacting with the ribosome, reflecting the ∼70Å translation of RNAP relative to the ribosome in TTC-LC relative to TTC-B (Figs. 2d-h, 3d-h). Coupling occurs solely, or nearly solely, through the coupling factor NusG or RfaH and the second coupling factor NusA (Figs. 2e-f, 3e-f). We refer to this new structural class as “TTC-LC”, where “LC” refers both to long-range coupling and loose coupling, reflecting the association of this structural class with long mRNA spacers between ribosome and RNAP, and the absence or near-absence of direct interaction between RNAP and ribosome.

For NusG-coupled, NusA-co-coupled TTCs containing an mRNA spacer length of 12 codons, we obtained both TTC-B and TTC-LC (Figs. 2-3; Extended Data Fig. 1; Extended Data Table 1). For NusG-coupled, NusA-co-coupled TTCs containing an mRNA spacer of 13 codons, and for RfaH-coupled, NusA-co-coupled TTCs containing mRNA spacers of length of 12 or 13 codons, we obtained only TTC-LC (Figs. 2-3; Extended Data Figs. 2-3; Extended Data Tables 1-2). Based on our previous results analyzing structures of TTCs with shorter mRNA spacers and on these new results with 12 or 13 codon mRNA spacers, we infer that, as anticipated, TTC-B only can accommodate, and be formed with, mRNA spacers up to 11-12 codons in length, and we infer that the new structural class, TTC-LC, is formed for mRNA spacers ≥12 codons in length. Structures of TTC-LC for mRNA spacers of 12 codons and mRNA spacers of 13 codons are superimposable (Figs 2c, 3c). The difference sin mRNA spacer length are accommodated by different extents of compaction and disorder of mRNA inside the RNAP RNA-exit channel and in mRNA in the “gap” between the RNAP RNA exit-channel and mRNA associated with the ribosome (Figs. 2e-f, 3e-f). Structures of the NusG-coupled, NusA-co-coupled TTC-LC and RfaH-coupled, NusA-co-coupled TTC-LC are superimposable (Figs. 2a, 3a).

Reflecting the difference in orientations of RNAP relative to the ribosome in TTC-B and TTC-LC, there are major differences in the interface between TEC and the ribosome in TTC-B and TTC-LC (Figs. 2e-h, 3e-h). In contrast to TTC-B, where the RNAP β’ zinc binding domain (β’ ZBD) makes direct protein-protein interaction with ribosomal protein S3 in the ribosome 30S subunit (314 Å^2^ buried surface area; Figs. 2g-h., 3g-h)^32–35^, in TTC-LC, RNAP makes no interactions with ribosomal protein S3 and makes no or almost no interactions with any other part of the ribosome, making at most contacts through 1-3 amino acids of the RNAP β’ zinc binding domain (ZBD) with ribosomal protein S10 (0-56 Å^2^; buried surface area; Figs. 2e-h, 3e-h).

In both TTC-B and TTC-LC, coupling factor NusG or RfaH bridges RNAP and the ribosome, with NusG-N or RfaH-N interacting with RNAP and NusG-C or RfaH-C interacting with ribosomal protein S10 in the ribosome 30S subunit (Figs. 2a, 2e-f, 3a, 3e-f). However, reflecting the difference in orientations of RNAP relative to the ribosome in TTC-B and TTC-LC, there is a major difference in the orientation of of NusG-N or RfaH-N relative to NusG-C or RfaH-C in TTC-B and TTC-LC (Figs. 2e-h, 3e-h). The difference in orientation is accommodated by a change in the conformation of the NusG or RfaH interdomain linker (Figs. 2e-h, 3e-h).

In both TTC-B and TTC-LC, coupling factor NusA makes a second bridge between RNAP and the ribosome (Figs. 1a-b, 2a, 2e-h, 3a, 3e-h). However, reflecting the difference in orientations of RNAP relative to the ribosome in TTC-B and TTC-LC, there are major differences in the bridging by NusA in TTC-B and TTC-LC (Figs. 2e-h, 3e-h). In TTC-B, the bridging by NusA involves extensive protein-protein interaction between NusA and ribosomal proteins S2 and S5 in the ribosome 30S subunit (Figs. 2g-h, 3g-h)^33–35^; in contrast, in TTC-LC, the bridging involves no protein-protein interactions between NusA and ribosomal proteins and is only indirect, involving only interactions between NusA and one face of the mRNA segment that, through its opposite face, interacts with ribosomal protein S3 in the ribosome 30S subunit (Figs. 2e-f, 3e-f).

In contrast to TTC-B, where mRNA proceeds directly from the RNAP RNA-exit channel to the RNA-helicase region^37^ of ribosomal protein S3 in the ribosome 30S subunit^32–35^ (Figs. 2g-h, 3g-h), in TTC-LC, there is a ∼70 Å gap between mRNA in the RNAP RNA-exit channel and mRNA interacting with the RNA-helicase region^37^ of ribosomal protein S3 in the ribosome 30S subunit (Figs. 2e-f, 3e-f, 4a-b). In our structures of TTC-LC, this mRNA segment is disordered (Figs. 2e-f, 3e-f, 4a-b); nevertheless, molecular modeling indicates that this mRNA segment makes no interactions with RNAP, coupling factors, or the ribosome, and indicate that this mRNA segment occupies a ∼45 Å wide, ∼70 Å long, solvent channel providing accessibility to bulk solvent outside TTC-LC (Fig 4a-b).

**Fig. 4.**
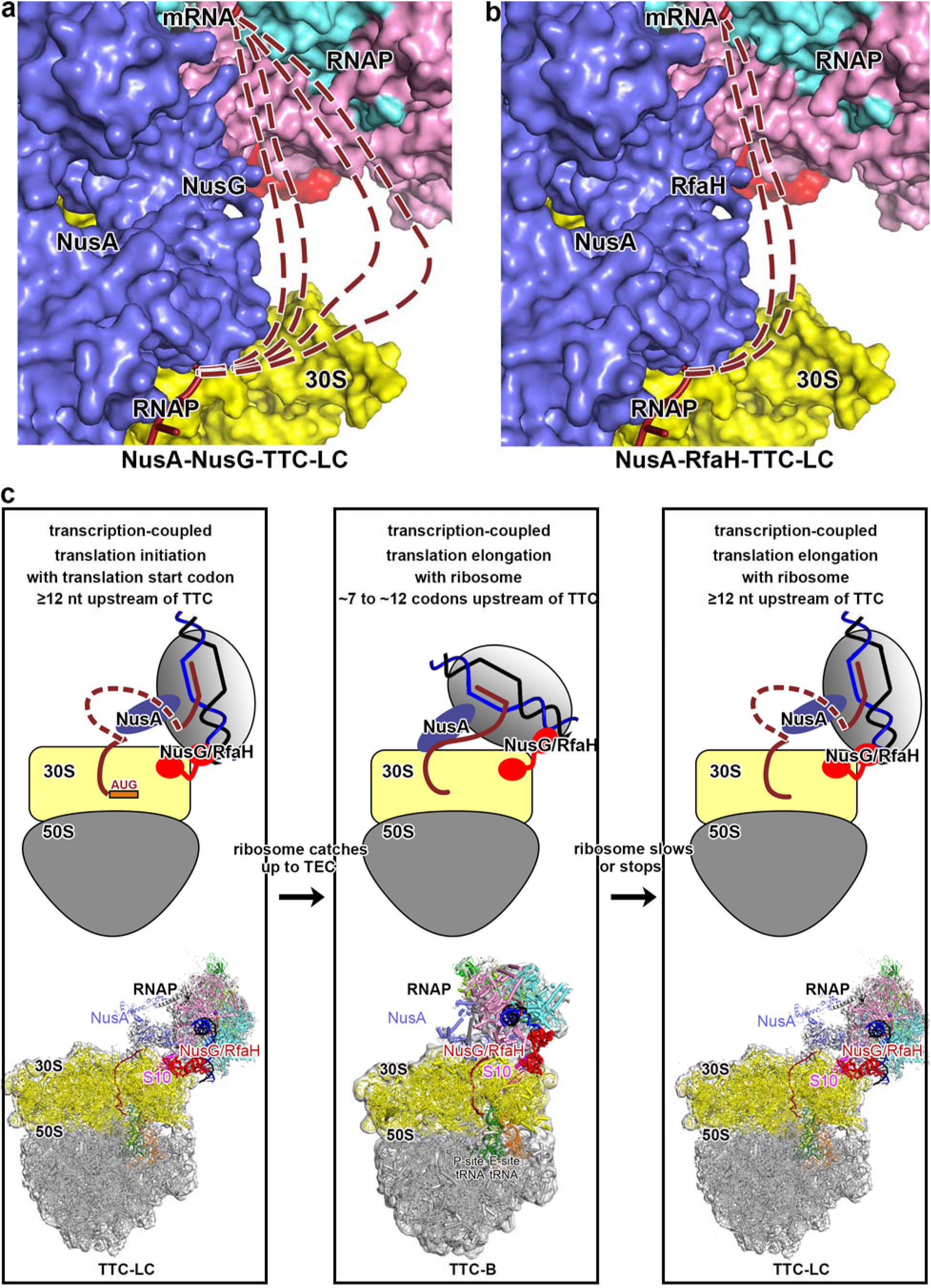
Accommodation of long mRNA spacers in TTC-LC through mRNA looping. **(a)** Inferred paths of mRNA spacers in NusA-NusG-TTC-LC. An mRNA spacer of 12 codons is accommodated without RNA looping (first brick-red dashed line, reading from left to right). mRNA spacers of 13, 17, and 20 codons (second, third, and fourth brick-red dashed lines, reading from left to right) are accommodated by progressively increasing extents of RNA looping into ∼45 Å wide, ∼70 Å long, solvent channel providing access to bulk solvent on exterior of TTC-LC (white space at center-right). Image shows view orientation that places segment of mRNA spacer in gap between mRNA in the RNAP RNA-exit channel (undashed brick-red ribbon at top) and mRNA interacting with ribosomal protein S3 in ribosome 30S subunit (solid brick-red ribbon at bottom) in plane of the image. Ribosome 30S subunit, yellow; other colors as in Figs. 1, 2a, and 3a. **(b)** as (a), but for NusA-RfaH-TTC-LC. **(c)** Proposal: TTC-LC serves as functional intermediate in assembling and disassembling TTC-B. Proposal has three steps (left, center, and right subpanels). In step 1 (left subpanel), TTC-LC mediates transcription-coupled translation initiation when mRNA spacer between RNAP and translation start codon is longer than maximum-length spacer compatible with TTC-B (n ≥ 12), and performs pre-TTC-B loose transcription-translation coupling. Subpanel shows schematic representation of TTC-LC, with looped out mRNA spacer as brick-red dashed line at center, and shows structure of TTC-LC at bottom. In step 2 (center subpanel), after ribosome “catches up” with RNAP, complex undergoes transition from TTC-LC to TTC-B, with no looping out of mRNA spacer, and mediates tight transcription-translation coupling for most transcription and translation. In step 3 (right subpanel), when ribosome stops moving and RNAP continues moving, resulting in increase in mRNA spacer length beyond maximum-length spacer compatible with TTC-B, the complex undergoes transition from TTC-B to TTC-LC, with looped out mRNA spacer (brick-red dashed line), and performs post-TTC-B loose transcription-translation coupling as TTC-LC.

We also have sought cryo-EM structures of NusG- and RfaH-coupled complexes containing mRNA spacers of 12 and 13 codons in the absence of the co-coupling factor NusA (Extended Data Figs. 4-5). Maps obtained in the absence of NusA show unequivocal density for RNAP interacting with the ribosome, and the density occupies a position that is superimposable on the position of density for TTC-LC obtained in the presence of NusA (Extended Data Figs. 4e-f, 5e-f). In the absence of NusA, map quality for RNAP is insufficient to define a unique fit and atomic model for RNAP interacting with the ribosome (Extended Data Figs. 4e, 5e), but density is decidedly clearer and decidedly more focused than density for “uncoupled TTCs” formed in the absence of NusG or RfaH (see ref. 31). We suggest that NusG- and RfaH-coupled TTC-LC can be formed both in the presence and in the absence of co-coupling factor NusA, but that the conformational heterogeneity and flexibility of the coupling is higher in the absence of NusA than in the presence.

### TTC-LC: structures of NusG-coupled TTCs containing very long--17 or 20 codons--mRNA spacers between RNAP and ribosome

In both NusG- and RfaH-coupled TTC-LC, the mRNA segment in the ∼70 Å gap between the mRNA in the RNAP RNA-exit channel and mRNA interacting with the RNA-helicase region of ribosomal protein S3 in the ribosome appears not to be engaged in protein-RNA interactions and occupies a ∼45 Å wide, ∼70 Å long solvent channel providing accessibility to bulk solvent the outside TTC-LC (Figs. 2e-f, 3e-f, 4-b). These observations suggest there likely are no constraints on the length or path of this mRNA segment and, thus, that longer mRNA spacers potentially can be accommodated through mRNA looping in this mRNA segment.

To test this hypothesis, in further studies, we determined structures of NusG-containing, NusA-containing TTCs having very long--17 codon and 20 codon (54 nt and 60 nt)--mRNA spacers (Fig. 2b). A 17 codon mRNA spacer exceeds the maximum-length mRNA spacer that can be accommodated solely through compaction and disorder in the RNAP RNA-exit channel of TTC-LC without mRNA looping, and a 20 codon mRNA spacer substantially exceeds the maximum-length spacer that could be accommodated solely through compaction and disorder in the RNAP RNA-exit channel without mRNA looping.

We obtained structures of NusG-coupled, NusA-co-coupled TTC-LC having 17 codon and 20 codon mRNA spacers (Figs. 2c, 4a-b; Extended Data Fig. 2d-e; Extended Data Table 1). The structures were superimposable on structures of NusG-coupled, NusA-co-coupled TTC-LC having 12 codon and 13 codon mRNA spacers (Extended Data Fig. 2f). As in the structures having 12 codon and 13 codon mRNA spacers,, mRNA in the RNAP RNA-exit channel and in the ∼70 Å gap between mRNA in the RNAP RNA-exit channel and mRNA interacting with the ribosome was disordered (Fig. 4a-b). Nevertheless, because a 17 codon mRNA spacer exceeds the maximum length of mRNA spacer that can be accommodated through compaction and disorder of mRNA in the RNAP RNA-exit channel, and because a 20 codon mRNA spacer substantially exceeds the maximum length of mRNA spacer that can be accommodated through compaction and disorder of mRNA in the RNAP RNA-exit channel, we conclude that mRNA in the ∼70 Å gap in the structures with 17 codon and 20 codon mRNA spacers must be accommodated through mRNA looping (Fig. 4a.b).

### TTC-LC transforms into TTC-B upon ribosome catch-up

We hypothesize TTC-LC serves as a functional precursor in forming TTC-B (Fig. 4c, left and center). We hypothesize that TTC-LC mediates transcription-coupled translation initiation when the mRNA spacer between RNAP and the translation start codon is longer than the maximum spacer length compatible with formation of TTC-B--which occurs frequently, in view of the relatively late loading of NusG and RfaH onto RNAP^9, 38–39^--and then mediates pre-TTC-B transcription-translation coupling, until the ribosome “catches up” to RNAP and forms TTC-B (Fig, 4, left and center).

To test the hypothesis that TTC-LC can transform into TTC-B when the ribosome advances faster than RNAP, decreasing the mRNA spacer length between RNAP and ribosome below the maximum spacer length compatible with TTC-B (”ribosome catch-up”), we prepared NusA-NusG-TTC-LC containing fMet-tRNA^fMet^ on a nucleic-acid scaffold having a 13-codon mRNA spacer (Fig. 5a, top); we then added Phe-tRNA^Phe^, EF-Tu, EF-G and GTP to “walk” the ribosome by 4 codons and thereby obtain a 9-codon mRNA spacer (Fig. 5a, bottom); and we then performed cryo-EM structure determination of the resulting complex (Fig. 5b-c; Extended Data Fig. 6; Extended Data Table 2). The structure obtained was NusA-NusG-TTC-B (subclass B2; Fig. 5b-c). We conclude that TTC-LC transforms into TTC-B upon ribosome catch-up.

**Fig. 5.**
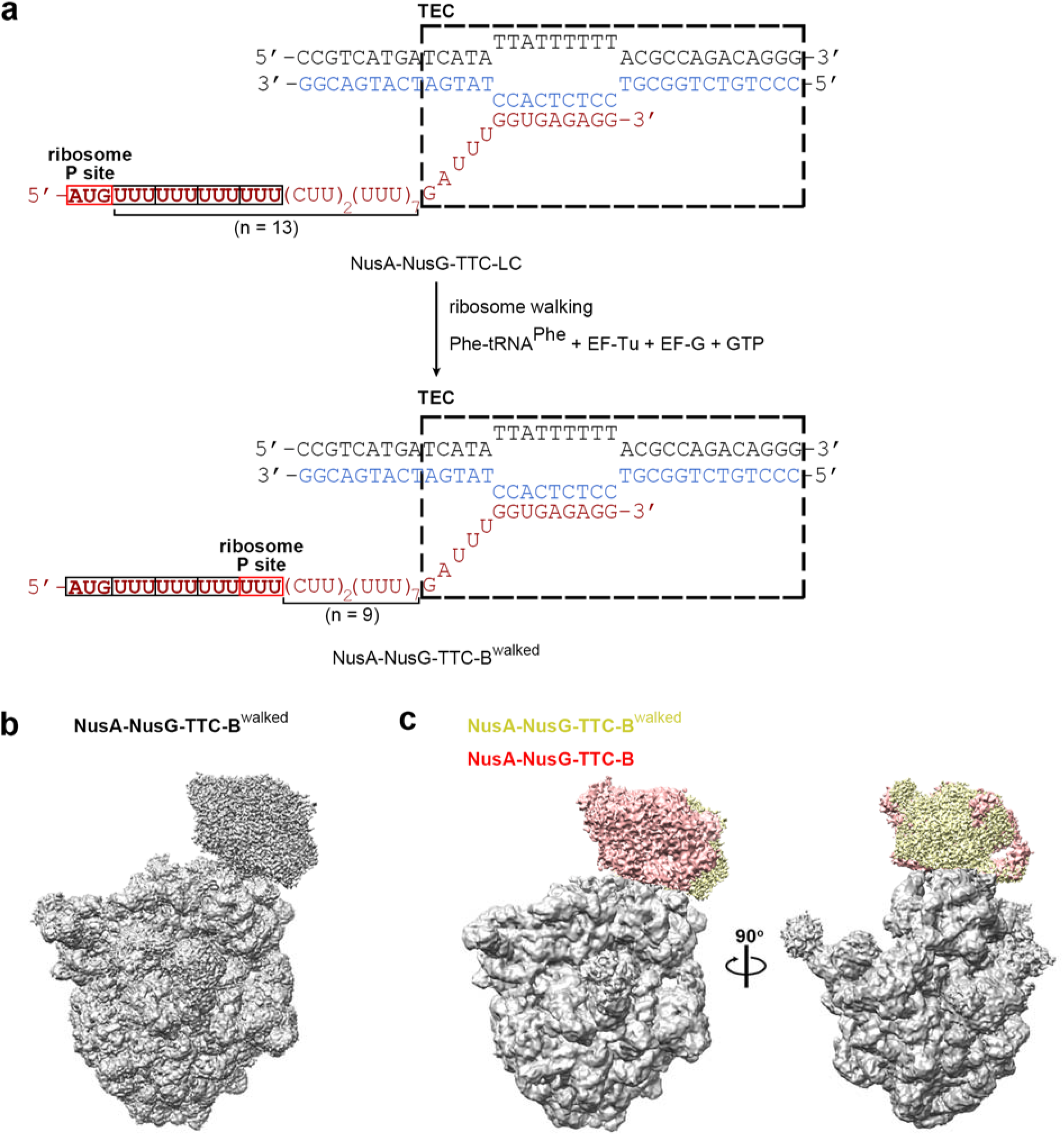
TTC-LC transforms into TTC-B upon ribosome catch-up. **(a)** Summary of experiment. Top, initial complex [NusA-NusG-TTC-LC (n = 13)]. Center, ribosome “walking.” Bottom, final complex [NusA-NusG-TTC-B^walked^ (n = 9)]. Codons translated during ribosome “walking,” bold, (CUU)_2_ halt site for ribosome “walking,” red. Other colors and features as in Fig. 1. **(b)** Superimposition of NusA-NusG-TTC-B^walked^ (TEC in yellow and ribosome in grey) on NusA-NusG- TTC-B^33^ (subclass B1; TEC in pink and ribosome in grey).

### TTC-B transforms into TTC-LC upon RNAP run-ahead

Similarly, we hypothesize that TTC-LC serves as a functional intermediate in disassembling TTC-B (Fig. 4c, center and right). We hypothesize TTC-LC mediates “post-TTC-B” transcription-translation coupling when the ribosome stops or slows but RNAP continues advancing, resulting in an mRNA spacer length longer than the maximum spacer length compatible with TTC-B (Fig 4c, right).

To test the hypothesis that TTC-B can transform into TTC-LC when RNAP advances faster than the ribosome, increasing the mRNA spacer length above the maximum spacer length compatible with TTC-B (”RNAP run-ahead”), we prepared NusA-NusG-TTC-B on a nucleic-acid scaffold having a 9-codon mRNA spacer (Fig. 6a, top); we then added CTP, GTP, and UTP to “walk” RNAP by 12 nt (4 codons) to obtain a 13-codon mRNA spacer (Fig. 6a, bottom); and we then performed cryo-EM structure determination of the resulting complex (Fig. 6b-c; Extended Data Fig. 7; Extended Data Table 2). The structure obtained was NusA-NusG-TTC-LC (Fig. 6b-c). We conclude that TTC-B transforms into TTC-LC upon RNAP run-ahead.

**Fig. 6.**
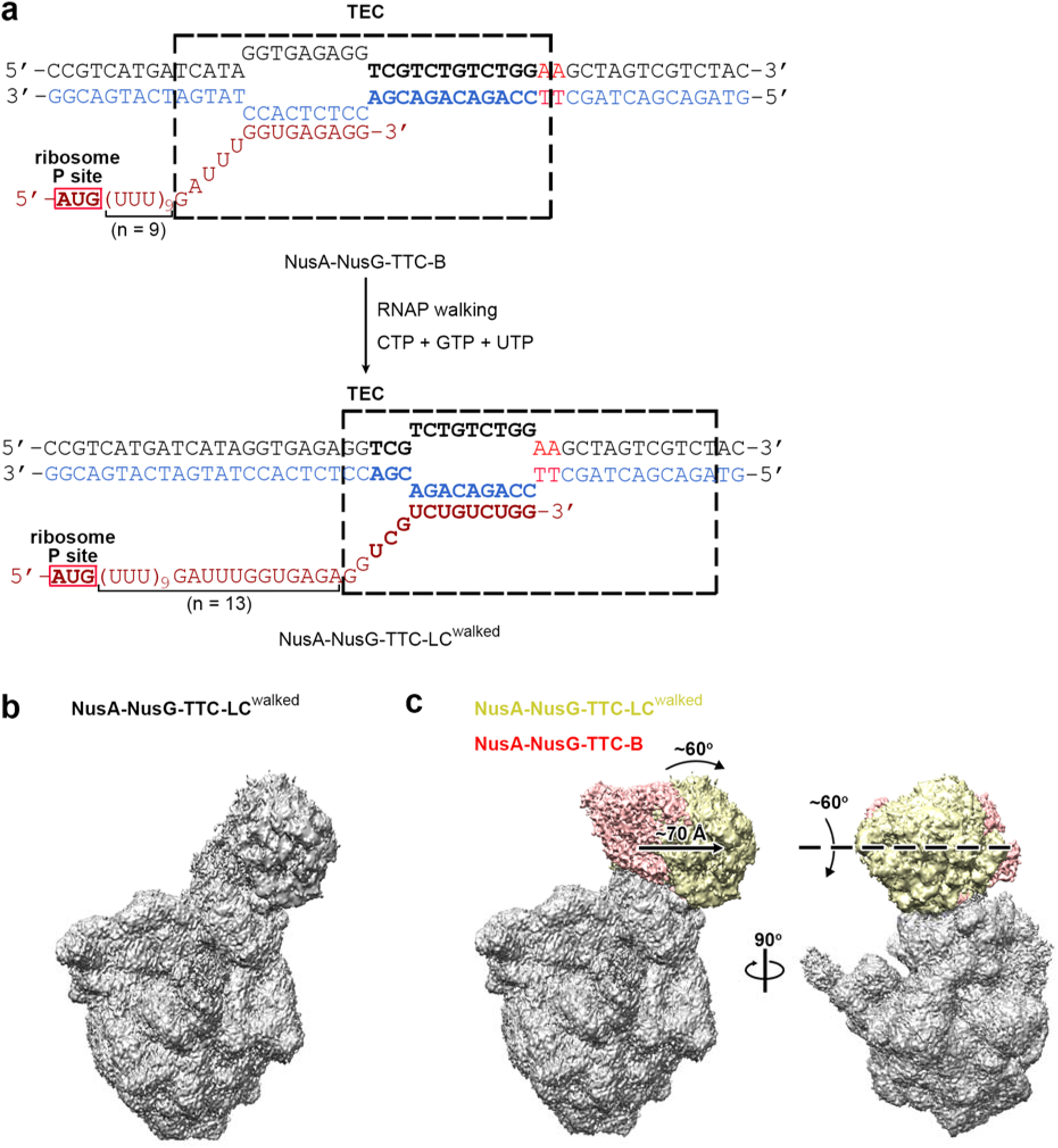
TTC-B transforms into TTC-LC upon RNAP run-ahead. **(a)** Summary of experiment. Top, initial complex [NusA-NusG-TTC-B (n = 9)]. Center, RNAP “walking.” Bottom, final complex [NusA-NusG-TTC-LC^walked^ (n = 13)]. Nucleotides transcribed during RNAP “walking,” bold, (A:T)_2_ halt site for RNAP “walking,” red. Other colors and features as in Fig. 1. **(b)** Superimposition of NusA-NusG-TTC-LC^walked^ (TEC in yellow and ribosome in grey) on NusA-NusG- TTC-B^33^ (subclass B1; TEC in pink and ribosome in grey).

### TTC-B, but not TTC-LC, is severely defective in hairpin-dependent intrinsic transcription termination

The structural differences between TTC-B and TTC-LC predict functional differences between TTC-B and TTC-LC in susceptibility to RNA-hairpin-dependent pausing and RNA-hairpin-dependent intrinsic termination. Molecular modelling indicates that TTC-B is unable to accommodate a pause hairpin^40–41^ or a termination hairpin^42^ without interaction of the hairpin with--and disruption of the hairpin by--the RNA-helicase region of ribosomal protein S3^32–35^. In contrast, molecular modeling indicates that TTC-LC is able to accommodate a pause or termination hairpin without interaction of the hairpin with the RNA-helicase region of ribosomal protein S3, by accommodating the hairpin within the ∼70 Å gap between RNAP and the RNA-helicase region of ribosomal protein S3 (Extended Data Fig. 8a). We therefore hypothesize that TTC-B, but not TTC-LC, is incompatible with hairpin-dependent transcription pausing and hairpin-dependent intrinsic transcription termination.

To test the hypothesis that TTC-B, but not TTC-LC, is incompatible with hairpin-dependent intrinsic transcription termination, we assessed the abilities of NusA-NusG-TTC-B, and NusA-NusG-TTC-LC to undergo termination at the tR2 terminator (Fig. 7). We prepared NusA-NusG-TEC and NusA-NusG-TTC-B positioned immediately upstream of the release site of the tR2 terminator (n = 9; Fig. 7a, top), and we prepared NusA-NusG-TEC and NusA-NusG-TTC-LC positioned immediately upstream of the release site of the tR2 terminator (n = 17; Fig. 7a, bottom). We then added ATP, CTP, GTP, and UTP to re-start transcription and assess termination at the tR2 terminator (detected as loss of ability to extend RNA past the release site of the tR2 terminator). The results show that NusA-NusG-TTC-B is >2-fold defective in termination at tR2, but that NusA-NusG-TTC-LC is only slightly defective in termination at tR2 (Fig. 7b-c), We conclude that TTC-B, but not TTC-LC, is severely defective in hairpin-dependent intrinsic transcription termination.

**Fig. 7.**
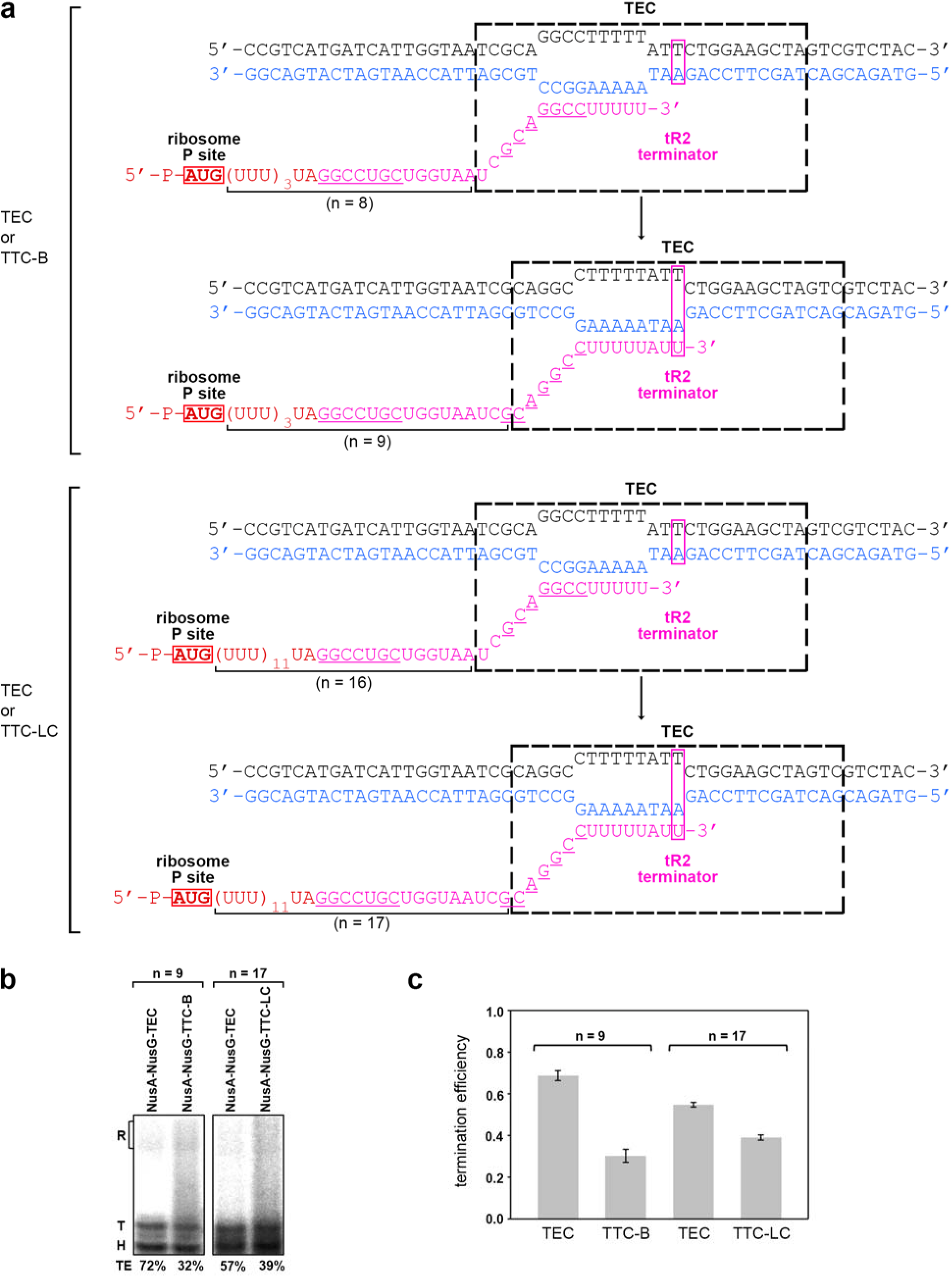
TTC-B, but not TTC-LC, is incompatible with hairpin-dependent intrinsic transcription termination. **(a)** Nucleic-acid scaffolds. Top, nucleic-acid scaffold enabling analysis of intrinsic termination by TEC or TTC-B (n =9). Bottom, nucleic-acid scaffold enabling analysis of intrinsic termination by TEC or TTC-LC (n =17). In each panel, upper subpanel shows scaffold upon “walking” of TEC to position 3 nt upstream of tR2 terminator release site (where at which position, ribosome and tRNA^fMet^ were added to allow formation of TTC), and lower subpanel shows scaffold upon further “walking” of TEC to tR2 terminator release site. Magenta, tR2 terminator; magenta underlining, tR2 hairpin-forming sequences; magenta rectangle, tR2 release site. Other colors and features as in Fig. 1. **(b)** RNA-extension data assessing intrinsic termination by TEC and TTC-B (left) and by TEC and TTC-LC (right). Termination is detected as loss of ability to extend RNA past the tR2 release site. H, RNA products in complexes halted 3 nt upstream of tR2 terminator release site, T, termination RNA products; R, run-off RNA products; TE, termination efficiencies. **(c)** Summary of termination efficiencies (mean+SEM; three independent determinations).

### Both TTC-B and TTC-LC are severely defective in Rho-dependent transcription termination

Molecular modeling indicates that both TTC-B and TTC-LC are incompatible with Rho-dependent transcription termination (Extended Data Fig. 8b).

To test the hypothesis that both TTC-B and TTC-LC are incompatible with Rho-dependent transcription termination, we assessed the abilities of NusA-NusG-TTC-B, NusA-NusG-TTC-LC, and NusA-NusG-TEC to undergo Rho-dependent termination (Fig. 8). We prepared NusA-NusG-TEC and NusA-NusG-TTC-B (n = 9; Fig. 8a, top), and we prepared NusA-NusG-TEC and NusA-NusG-TTC-LC (n = 17; Fig. 8a, bottom). We then added Rho, Rho primary binding site (PBS) ligand dC75^24^, and dATP to enable Rho-dependent termination, and we then added ATP, CTP, GTP, and UTP to re-start transcription and assess Rho-dependent termination (detected as loss of ability to extend RNA). The results show that both NusA-NusG-TTC-B and NusA-NusG-TTC-LC are >2-fold defective in Rho-dependent termination (Fig. 8b-c). We conclude that both TTC-B and TTC-LC are severely defective in Rho-dependent transcription termination.

**Fig. 8.**
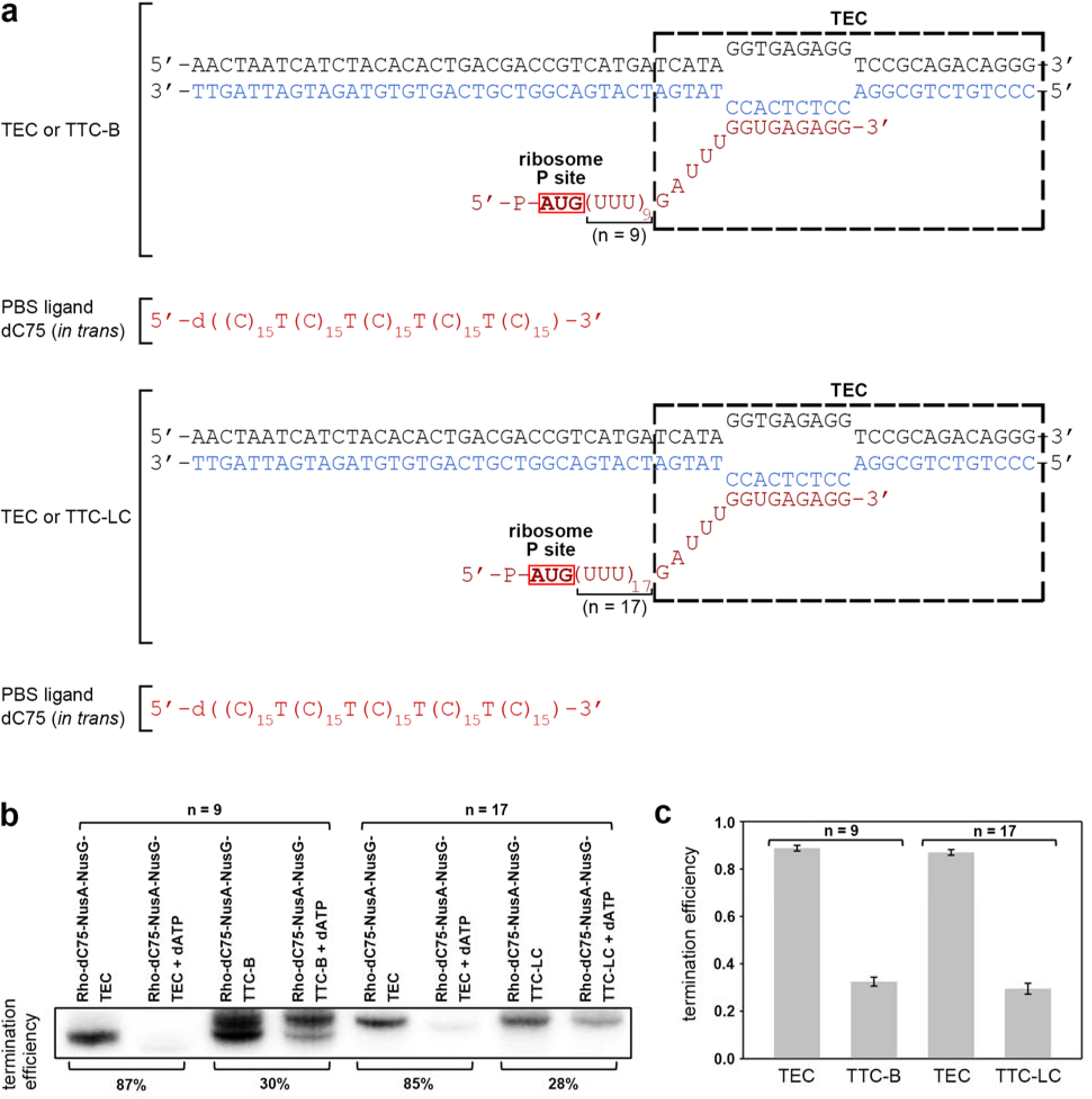
Both TTC-B and TTC-LC are incompatible with Rho-dependent transcription termination. **(a)** Nucleic-acid scaffolds. Top, nucleic-acid scaffold enabling assembly and analysis of TEC or TTC-B (n = 9). Bottom, nucleic-acid scaffold enabling assembly and analysis of TEC or TTC-LC RNA (n = 17). Colors and features as in Fig. 1. **(b)** RNA-extension data assessing Rho-dependent termination by TEC and TTC-B (left) and by TEC and TTC-LC (right). Termination is detected as loss of ability to extend RNA. Termination efficiencies are at bottom.

## Discussion

Our results define a new structural class of NusG- and RfaH-coupled TTCs (Figs. 2–4). This new class accommodates mRNA spacers between RNAP and the ribosome that are longer than those that can be accommodated by the previously defined structural class TTC-B. In this new structural class, there is no or almost no direct interaction between RNAP and the ribosome. Coupling is mediated by bridging of RNAP and the ribosome through the coupling factor NusG or RfaH and by supplementary indirect bridging of RNAP and the ribosome through the coupling factor NusA. In this new structural class, there is a ∼70 Å gap between mRNA in the RNAP RNA-exit channel and mRNA interacting with the ribosome 30S subunit. mRNA in the ∼70 Å gap appears to make no protein-RNA interactions and to be accessible to the exterior of TTC-LC through a ∼45 Å wide, ∼70 Å long, solvent channel. In this new structural class, mRNA in the ∼70 Å gap can accommodate very long, potentially unlimited-lengh, mRNA spacers through RNA looping. We refer to this new class as “TTC-LC.” where “LC” denotes both long-range coupling, reflecting the fact that this class is associated with long mRNA spacers, and loose coupling, reflecting the absence or near absence of direct RNAP-ribosome interaction. Our results further show that TTC-LC can transform into TTC-B upon ribosome catch-up (Fig. 5) and that TTC-B can transform into TTC-LC upon RNAP run-ahead (Fig. 6), Finally, our results show that TTC-B, but not TTC-LC, is severely defective in hairpin-dependent intrinsic transcription termination (Fig. 7), and that both TTC-B and TTC-LC are severely defective in Rho-dependent transcription termination (Fig. 8),

This new structural state, TTC-LC, explains, and accounts for, the single-molecule fluorescence-resonance energy-transfer observations of Qureshi and Duss^36^. Specifically, TTC-LC enables long-range coupling with very long mRNA spacers between RNAP and the ribosome, involves coupling through NusG or RfaH and NusA, and involves, due to the absence or near absence of RNAP-ribosome interaction, a lower stability of coupling^36^.

This new structural state, TTC-LC, also explains, and accounts for, the biochemical observations of N et al.^43^, who presented evidence that TTCs having mRNA spacers <10 codons in length, but not TTCs having longer mRNA spacers, are defective in intrinsic transcription termination in the *E. coli gal* operon^43^.

We propose that TTC-LC is a functional coupling complex and serves as a functional intermediate in forming and disassembling TTC-B (Fig. 4c). We propose that TTC-LC mediates transcription-coupled translation initiation when the mRNA spacer between RNAP and the translation start codon is longer than the maximum spacer length compatible with formation of TTC-B--which occurs frequently, in view of the relatively late loading of NusG and RfaH onto RNAP^9, 38–39^--and then mediates pre-TTC-B transcription-translation coupling, until the ribosome “catches up” to RNAP and forms TTC-B (Fig, 4, left and center). We also hypothesize that TTC-LC mediates “post-TTC-B” transcription-translation coupling when the ribosome stops moving but RNAP continues moving, resulting in an mRNA spacer length longer than the maximum spacer length compatible with TTC-B (Fig 4, right). The observation that NusG and RfaH often load onto RNAP at positions downstream of the translation start codon that would yield mRNA spacers between RNAP and the ribosome substantially longer than those that can be accommodated in TTC-B^9, 38–39^ has posed a challenge to the hypothesis that TTC-B functionally mediates transcription-translation coupling^9, 12–13^. The discovery of this new structural class, TTC-LC, able to mediate transcription-translation coupling with substantially longer mRNA spacers, resolves this challenge.

Key priorities for future work include demonstration that both TTC-B and TTC-LC are present in cells, by use of *in situ* structural biology through cryo-electron tomography^44^, and assessment of the effects of TTC-LC on transcription pausing and transcription termination, by use of the single-molecule fluorescence resonance energy transfer assays of Qureshi and Duss^36^.

## Acknowledgments

We thank the Rutgers CryoEM and Nanoimaging Facility (supported by NIH grant S10OD036338), the Stanford-SLAC Cryo-EM Center (supported by NIH grant GM129541), and the Cryo-EM Center of the Shanghai Institute of Immunology and Infection (supported by National Natural Science Foundation grant 32521004 to C.W.) for microscope access. We thank Emre Firlar for technical assistance with cryo-EM data collection.

## Author contributions

C.W., V.M., K.K., and R.H.E. designed experiments. V.M., G.B., and K.K. prepared proteins and nucleic acids. V.M, C.W., S.S., L.Y., J.Z., and J.T.K. performed cryo-EM data collection. K.K. performed termination assays. C.W., V.M., S.S., L.Y., K.K., J.Z., B.E.N., and R.H.E. analyzed data. C.W., V.W., S.S., K.K.,, J.Z., and R.H.E. prepared figures. R.H.E. wrote the manuscript.

## Data availability statement

Cryo-EM maps have been deposited in the Electron Microscopy Database (EMDB accession codes 43335, 43387, 43388, 43389, 43390, 43391, 65509, 72553, 72554, and 72646, and atomic coordinates have been deposited in the Protein Database (PDB accession codes 8VL1, 8VOO, 8VOP, 8VOQ, 8VOR, 8VOS, 9W0N, and 9Y79). Unique biological materials will be made available to qualified investigators on request.

## Methods

### *E. coli* RNAP core enzyme, NusA, NusG, RfaH, Rho, and 70S ribosomes

Hexahistidine-tagged *E. coli* RNAP core enzyme, *E. coli* NusA, *E. coli* NusG, *E. coli* Rho, and *E. coli* 70S ribosomes were prepared as described ^24, 33^. *E. coli* Ef-Tu, *E. coli* Ef-G, *E. coli* fMet-^tRNAfMet^, and *E. coli* Phe-tRNA^Phe^ were prepared as described^35, 45, 46^. *E. coli* RfaH^E48A^, an RfaH derivative that contains a substitution in RfaH-N that disrupts interactions between RfaH-N and RfaH-C that stabilize an inactive conformational state of RfaH^5^, was prepared as described^34^.

### *E. coli* tRNA^fMet^

*E. coli* tRNA^fMet^ was purchased (MP Biomedical), dissolved to 100 μM in 5 mM Tris-HCl, pH 7.5, and stored in aliquots at -80°C.

### Nucleic acids

Oligodeoxyribonucleotides and oligoribonucleotides (sequences in Figs. 2b, 3b, 5a, 6a, 7a, and 8a) were purchased (Integrated DNA Technologies), PAGE-purified, dissolved in annealing buffer (5 mM Tris-HCl, pH 7.5) to 1 mM, and stored at -80°C in aliquots.

Nucleic-acid scaffolds (sequences in Figs. 2b, 3b, 5a, 6a, 7a, and 8a) were prepared as described^33–34^, mixing 60 mM nontemplate-strand oligodeoxyribonucleotide, 60 mM template-strand oligodeoxyribonucleotide, and 60 mM oligoribonucleotide in 100 µl annealing buffer (5 mM Tris-HCl, pH 7.5), heating 10 min at 95°C, and cooling slowly (3 h) to 22°, and nucleic-acid scaffolds were stored in aliquots at -80°C. Partial nucleic-acid scaffolds comprising template-strand oligodeoxyribonucleotide and oligoribonucleotide were prepared in the same manner, but omitting nontemplate-strand oligodeoxyribonucleotide.

### Termination assays: intrinsic termination

Intrinsic termination was assayed in RNA-extension assays, using nucleic-acid scaffolds that enabled preparation and analysis of TEC or TTC-B containing 5’-^32^P-labelled RNA and positioned immediately upstream of the release site for the tR2 terminator (n = 9; Fig. 7a, top) or preparation and analysis of TEC or TTC-LC containing 5’-^32^P-labelled RNA and positioned immediately upstream of the release site for the tR2 terminator (n = 17; Fig. 7a, bottom). TECs and TTCs were assembled on the scaffolds; then ATP, CTP, GTP, and UTP were added to re-start transcription and assess intrinsic termination (detected as loss of ability to extend RNA past the release site of the tR2 terminator).

Reaction mixtures containing 0.5 µM RNAP core enzyme immobilized on 5 μl Ni-NTA-agarose beads (QIAGEN)^47^ and 0.5 µM partial nucleic-acid scaffold comprising template-strand oligodeoxyribonucleotide and oligoribonucleotide 5’-^32^P-AUGUUUUUUUUUUAGGCCUGCUGGUAAUCG-3’ or 5’-^32^P- AUGUUUUUUUUUUUUUUUUUUUUUUUUUUUUUUUUUUAGGCCUGCUGGUAAUCG -3’ (1.9 Bq/fmol) in 10 μl transcription buffer (50 mM Tris-HCl, pH 7.9, 10 mM MgCl_2_, 20 mM KCl, and 1 mM dithiothreitol) were incubated 10 min at 22°C; were supplemented with 1 μl 6 µM nontemplate-strand oligodeoxyribonucleotide and incubated 5 min at 22°C; were supplemented with 0.5 μl 500 μM ATP and 500 μM CTP (>99% purity; ThermoFisher) and incubated 5 min at 37°C; were centrifuged (15,000 x g) 1 min, washed three times with 50 μl transcription buffer, and re-suspended in 10 μl transcription buffer at 22°C; were supplemented with 0.5 μl 500 μM CTP, 500 μM GTP, and 500 μM UTP (>99% purity; ThermoFisher) and incubated 5 min at 37°C; were centrifuged (15,000 x g) 1 min, washed with 50 μl transcription buffer, and re-suspended in 10 μl transcription buffer at 22°C; were eluted from Ni-NTA-agarose beads by addition of 0.5 μl 2 M imidazole, incubation 5 min at 22°C, centrifugation (15,000 x g) 1 min, and transfer of supernatant to fresh tubes; were supplemented with 1 μl 30 μM NusA and 0.45 μl 68 μM NusG and incubated 5 min at 22°C; and were supplemented with 0.75 μl ribosome storage buffer [20 mM Tris-HCl, pH 7.5, 30 mM KCl, 20 mM Mg(OAc)_2_, and 4 mM 2-mercaptoethanol] and 1 μl water for TECs or 0.75 μl 32 μM 70S ribosomes in ribosome storage buffer and 1 μl 100 mM tRNA^fMet^ for TTCs and incubated 5 min at 22°C. The resulting reaction mixtures then were supplemented with 1.5 μl 1 mM ATP, 1 mM CTP, 1 mM GTP, and 1 mM UTP (>99% purity; ThermoFisher) and incubated 5 min at 37°C (to “chase” complexes); were mixed with 1 μl ice-cold 0.5 M EDTA to stop reactions, were phenol-extracted, and were ethanol-precipitated. Reaction products were resolved by electrophoresis on 7 M urea, 10% polyacrylamide gels (19:1 acrylamide:bis-acrylamide), were detected using storage-phosphor imaging (Typhoon PhosphorImager, GE Healthcare), and were quantified using ImageQuant 5.2 (GE Healthcare).

### Termination assays: Rho-dependent termination

Rho-dependent termination was assessed in RNA-extension scaffold assays^24^, using nucleic-acid scaffolds that direct assembly of TEC or TTC-B containing 5’-phosphoryated RNA (n = 9; Fig. 8a, top) or assembly of TEC or TTC-LC with 5’-phosphoryated RNA (n = 17; Fig. 8a, bottom). TECs and TTCs were assembled on the scaffolds; then Rho, Rho PBS ligand dC75^24^, and dATP were added to enable Rho-dependent termination; and then [α32P]ATP, CTP, GTP, and UTP were added to re-start transcription and assess Rho-dependent termination (detected as loss of ability to extend RNA).

Reaction mixtures containing 0.5 µM RNAP core enzyme and 0.5 µM partial nucleic-acid scaffold comprising template-strand oligodeoxyribonucleotide and 5’-phosphoryated oligoribonucleotide in 10 μl transcription buffer were incubated 10 min at 22°C; were supplemented with 1 μl 6 µM nontemplate-strand oligodeoxyribonucleotide and incubated 5 min at 22°C, were supplemented with 1 μl 30 μM NusA and 0.45 μl 68 µM NusG and incubated 5 min at 22°C, were supplemented with 0.75 μl ribosome storage buffer and 1 μl water for TECs or 0.75 μl 32 μM 70S ribosomes in ribosome storage buffer and 1 μl 100 mM tRNA^fMet^ for TTCs and incubated 5 min at 22°C, were supplemented with 1 μl 35 μM dC75 and incubated 1 min at 22°C, and were supplemented with 1 μl 30 µM Rho and incubated 5 min at 22°C. The resulting reaction mixtures then were supplemented with 0.75 μl 20 mM dATP in transcription buffer and incubated 2 min at 37°C (to enable termination), were supplemented with 1 μl 50 μM [α^32^P]ATP (3.5 Bq/fmol), 50 μM CTP, 50 µM GTP, and 50 µM UTP (>99% purity; ThermoFisher), and incubated 3 min at 37°C (to “chase” non-terminated complexes), and were mixed with 20 μl 2x stop buffer (95% formamide, 0.2 mg/ml heparin, 25 mM EDTA, 0.03% SDS, and 0.03% bromophenol blue) at 22°C. Reaction products were resolved by electrophoresis on 7 M urea, 22% polyacrylamide gels (19:1 acrylamide:bis-acrylamide), were detected using storage-phosphor imaging (Typhoon PhosphorImager, GE Healthcare), and were quantified using ImageQuant 5.2 (GE Healthcare).

### Cryo-EM structure determination: sample preparation

TTCs of Figs. 2–4 were prepared as described^33–34^, mixing 10 µl 20 µM RNAP core enzyme (in RNAP storage buffer: 10 mM Tris-HCl, pH 7.6, 100 mM NaCl, 0.1 mM EDTA, and 5 mM dithiothreitol), 0 or 30 μl 30 μM NusG or RfaH^E48A^ (in RNAP storage buffer), 0 or 16 μl 60 μM NusA (in RNAP storage buffer), 4 µl 60 µM nucleic-acid scaffold (in 5 mM Tris-HCl, pH 7.5), 10 μl 10x 70S ribosome storage buffer, and 26 μl (reactions without NusA) or 10 μl (reactions with NusA) water; incubating 10 min at 22°C; adding 10 µl 100 µM tRNA^fMet^ (in 5 mM Tris-HCl, pH 7.5), 5 µl 40 µM 70S ribosome (in 70S ribosome storage buffer), and 5 μl water; and further incubating 10 min at 22°C.

NusA-NusG-TTC-B^walked^ (n = 9) of Fig. 5 was prepared in two stages. In the first stage, NusA-NusG-TTC-LC (n = 13) was prepared by mixing 10 µl 20 µM RNAP core enzyme (in RNAP storage buffer), 30 μl 30 μM NusG (in RNAP storage buffer), 16 μl 60 μM NusA (in RNAP storage buffer) and 4 µl 60 µM nucleic-acid scaffold of Fig 5a (in annealing buffer), 10 μl 10x 70S ribosome storage buffer (10x 20 mM Tris-HCl, pH 7.5, 30 mM KCl, 20 mM Mg(OAc)_2_, and 4 mM 2-mercaptoethanol), and 18 μl water and incubating 10 min at 22°C;; and adding 2 µl 100 µM fMet-tRNA^fMet^ (in 5 mM Tris-HCl, pH 7.5), and 10 µl 20 µM 70S ribosomes (in 70S ribosome storage buffer); and further incubating 10 min at 37°C. In the second stage, the ribosome in NusA-NusG-TTC-LC (n = 13) was “walked” forward by 4 codons, yielding NusA-NusG-TTC-B^walked^ (n = 9), by addition of Phe-tRNA^Phe^/EF-Tu/GTP ternary complex (prepared by pre-incubating 10 µl 100 µM Phe-tRNA^Phe^, 12.5 µl 120 µM EF-Tu, and 7.5 µl 100 mM GTP 5 min at 4°C), 44.5 µl 360 µM EF-G, and 20 µl 100 mM GTP and incubation 10 min at 37°C.

NusA-NusG-TTC-LC^walked^ (n = 13) of Fig. 6 was prepared in two stages. In the first stage, NusA-NusG-TTC-B (n = 9) was prepared by mixing 10 µl 20 µM RNAP core enzyme (in RNAP storage buffer), 4 μl 60 μM partial nucleic-acid scaffold comprising template-strand oligodeoxyribonucleotide and oligoribonucleotide of Fig 6a (in annealing buffer), 10 μl 10x 70S ribosome storage buffer, and 8 μl water and incubating 10 min at 22°C; adding 2 μl 250 μM nontemplate-strand oligodeoxyribonucleotide of Fig. 6a (in annealing buffer) and further incubating 10 min at 22°C; adding 30 μl 30 μM NusG (in RNAP storage buffer) and 16 μl 60 μM NusA (in RNAP storage buffer) and further incubating 5 min at 22°C; and adding 10 µl 100 µM tRNA^fMet^ (in 5 mM Tris-HCl, pH 7.5), 5 µl 40 µM 70S ribosome (in 70S ribosome storage buffer), and 5 μl water; and further incubating 10 min at 22°C. In the second stage, the RNAP in NusA-NusG-TTC-B (n = 9) was “walked” forward by 4 codons, yielding NusA-NusG-TTC-LC^walked^ (n = 13), by addition of 3 μl 2 mM CTP, 2 mM GTP, and 2 mM ATP and incubation 10 min at 22°C.

In all cases, TTCs prepared as described above were transferred to pre-chilled 0.5 ml Amicon Ultracel 30K concentrators (EMD Millipore), concentrated to 35 µl by centrifugation (15 min at 20,000xg at 4°C), incubated 30 min on ice for 30 min, supplemented with 3.8 µl of ice-cold 80 mM CHAPSO, and immediately applied to grids.

EM grids were prepared as described^33–34^, using a Vitrobot Mark IV autoplunger (FEI/ThermoFisher), with the environmental chamber at 22°C and 100% relative humidity. Samples (3 μl) were applied to 2/1 Quantifoil Cu 300 holey-carbon grids [Quantifoil; glow-discharged 60 s using a PELCO glow-discharge system (Ted Pella)], grids were blotted with #595 filter paper (Ted Pella) for 7 s at 22°C, and grids were flash-frozen by plunging in a liquid ethane cooled with liquid N_2_, and grids were stored in liquid N_2_.

### Cryo-EM structure determination: data collection and data reduction

Cryo-EM data for NusA-NusG-TTC-B (n = 12) and NusA-NusG-TTC-LC (n = 12), NusA-NusG-TTC-LC (n = 13), NusA-NusG-TTC-LC (n = 17), NusA-NusG-TTC-LC (n = 20), and NusA-RfaH-TTC-LC (n = 12) were collected at the Rutgers University Cryo-EM and Nanoimaging Core Facility, using a 200 kV Talos Arctica (FEI/ThermoFisher) electron microscope equipped with a GIF Quantum K2 direct electron detector (Gatan) (Extended Data Figs. 1-3). Data were collected automatically in counting mode, using EPU (FEI/ThermoFisher), a nominal magnification of 130,000x, a calibrated pixel size of 1.038 Å/pixel, and a dose rate of 4.8 electrons/pixel/s. Movies were recorded at 200 ms/frame for 6 s (30 frames), resulting in a total radiation dose of 26.7 electrons/Å^2^. Defocus range varied between -1.25 µm and -2 µm. For NusA-NusG-TTC-B (n = 12) and NusA-NusG-TTC-LC (n = 12) combined, NusA-NusG-TTC-LC (n = 13), NusA-NusG-TTC-LC (n = 17), NusA-NusG-TTC-LC (n = 20), and NusA-RfaH-TTC-LC (n = 12), datasets of 4,692, 4,544, 4,381, and 5,017 micrographs, respectively, were recorded from single grids over 2-3 days.

Micrographs were gain-normalized and defect-corrected. Data were processed as summarized in Extended Data Figs. 1-3. Data processing was performed using a Tensor TS4 Linux GPU workstation with four GTX 1080 Ti graphic cards (NVIDIA). Dose-weighting motion correction (3x3 tiles; b-factor = 150) were performed using Motioncor 2^48^. Contrast-transfer-function (CTF) estimation was performed using CTFFIND-4.1^49^. Subsequent image processing was performed using Relion 3.0^50^. Automatic particle picking with Laplacian-of-Gaussian filtering yielded initial particle sets for NusA-NusG-TTC-B (n = 12) and NusA-NusG-TTC-LC (n = 12) combined, NusA-NusG-TTC-LC (n = 13), NusA-NusG-TTC-LC (n = 17), NusA-NusG-TTC-LC (n = 20), and NusA-RfaH-TTC-LC (n = 12) of 17,581, 3,213, 122,400, 209,053, and 90,290 particles, respectively. Particles were extracted into 500x500 pixel boxes and subjected to rounds of reference-free 2D classification and removal of poorly populated classes, yielding selected particle sets for NusA-NusG-TTC-B (n = 12), NusA-NusG-TTC-LC (n = 12), NusA-NusG-TTC-LC (n = 13), NusA-NusG-TTC-LC (n = 17), NusA-NusG-TTC-LC (n = 20), and NusA-RfaH-TTC-LC (n = 12) of 3,494, 6,645, 1,516, 24,109, 32,821, and 20,048 particles, respectively. The resulting selected particle sets were 3D-classified with C1 symmetry, using a *de novo* 3D template created using 3D_initial_model under Relion 3.0. For each complex, classes exhibiting strong, well-defined densities assignable to TEC and ribosome and having a spatial relationship of density for TEC and ribosome consistent with a TTC were combined and were 3D auto-refined using a mask with a diameter of 450 Å. The resulting 3D auto-refined particles were further refined using a soft mask and solvent flattening and were post-processed, yielding reconstructions for NusA-NusG-TTC-B (n = 12), NusA-NusG-TTC-LC (n = 12), NusA-NusG-TTC-LC (n = 13), NusA-NusG-TTC-LC (n = 17), NusA-NusG-TTC-LC (n = 20), and NusA-RfaH-TTC-LC (n = 12) at 6.2, 6.0, 6.2, 3.6, 3.8, and 5.3 Å overall resolution, respectively, as determined from gold-standard Fourier shell correlation (FSC; Extended Data Figs. 1d, 2d, and 3d; Extended Data Tables 1-2). The initial atomic models for TTCs were built as described^33–34^. Refinement of the initial models was performed as described^33–34^. The initial models were subjected to rounds of manual refinement using Coot^51^ and auto-refinement using Phenix real-space refinement^52^.

Cryo-EM data for NusA-RfaH-TTC-LC (n = 13) were collected at the Stanford-SLAC Cryo-EM Center, using a 300 kV Titan Krios (FEI/ThermoFisher) electron microscope equipped with a Gatan K3 Quantum direct electron detector (Gatan) (Extended Data Fig. 3). Data were collected automatically in counting mode using EPU (FEI/ThermoFisher), a nominal magnification of 81,000x, a calibrated pixel size of 0.86 Å/pixel, and a dose rate of 40 electrons/pixel/s. Movies were recorded at 30 ms/frame for 3 s (100 frames), resulting in a total radiation dose of 45 electrons/Å^2^. Defocus range varied between -0.8 µm and -2.5 µm. A dataset of 7,067 micrographs was recorded from one grid over 2 days. Data were processed as above, yielding a final set of 15,810 particles that produced a reconstruction of 3.4 Å overall resolution (Extended Data Fig. 3d; Extended Data Table 2).

Cryo-EM data for NusA-NusG-TTC-B^walked^ (n = 9; subclass B2) of Fig. 5 were collected at the Cryo-EM Center of Shanghai Institute of Immunity and Infection, using a 300 kV Titan Krios (FEI/ThermoFisher) electron microscope equipped with a Gatan K3 Quantum direct electron detector (Gatan) (Extended Data Table 2). Data were collected automatically in super-resolution mode using EPU (FEI/ThermoFisher), a nominal magnification of 105,000x, a calibrated pixel size of 0.824 Å/pixel, and a dose rate of 50 electrons/pixel/s. Movies were recorded at 50 ms/frame for 2 s (40 frames), resulting in a total radiation dose of 73 electrons/Å^2^. Defocus range varied between -1.0 µm and -2.2 µm. A dataset of 10,431 micrographs was recorded from one grid over one day. Data were processed as above, yielding a final set of 81,802 particles that produced a reconstruction of 3.1 Å overall resolution (Extended Data Fig. 6; Extended Data Table 2). Because local resolution for NusA-NusG-TEC was superior to that in our previous structure of NusA-NusG-TTC-B (n = 9; subclass B2)^33^, an atomic model was built and refined (procedures as described above).

Cryo-EM data for NusA-NusG-TTC-LC^walked^ (n = 13) of Fig. 6 were collected at the Rutgers University Cryo-EM and Nanoimaging Core Facility, using a 300 kV Titan Krios (FEI/ThermoFisher) electron microscope equipped with a Gatan K3 Quantum direct electron detector (Gatan) (Extended Data Table 2). Data were collected automatically in counting mode using EPU (FEI/ThermoFisher), a nominal magnification of 130,000x, a calibrated pixel size of 0.685 Å/pixel, and a dose rate of 9.16 electrons/pixel/s. Movies were recorded at 62.5 ms/frame for 2.5 s (40 frames), resulting in a total radiation dose of 50 electrons/Å2. Defocus range varied between -0.8 µm and -2.0 µm. A dataset of 12,587 micrographs was recorded from one grid over one day. Data processing was performed using cryoSPARC v4.2.1^53^. Following patch motion correction, patch CTF estimation, and manual inspection to remove micrographs with ice contamination or poor image quality, particles were picked from a random subset of 5,000 micrographs using cryoSPARC Blob Picker, 2D classification was performed to generate a 2D template, and particles were autopicked from 11,391 micrographs using the 2D template and cryoSPARC Template Picker, yielding an initial particle set of 592,698 particles. Particles were extracted into 128x128 pixel boxes (6x downscaled) and subjected to six rounds of reference-free 2D classification and two rounds of heterogeneous refinement, yielding a selected particle set of 122,143 particles. The selected particles were re-extracted into 512x512 pixel boxes, subjected to particle subtraction using a ribosome mask, and subjected to focussed 3D classification, yielding a selected subclass of 32,979 particles corresponding to NusA-NusG-TTC-LC^walked^. The selected-subclass particles were re-extracted into 512x512 pixel boxes (2x downscaled) and subjected to non-uniform refinement with global CTF refinement, yielding a reconstruction with global resolution of 3.20 Å as determined from gold-standard Fourier shell correlation (map 1; Extended Data Fig. 7a,d–e). To improve map quality for NusA-NusG-TEC, the selected-subclass particles were subjected to local refinement using a mask for NusA-NusG-TEC, yielding a locally refined map for NusA-NusG-TEC with a resolution of 8.50 Å as determined from gold-standard Fourier shell correlation (map 1a; Extended Data Fig. 7a, d-e). The global map (map 1) and the locally refined map (map 1a) were combined in Chimera^54^ using the “vop maximum” command to generate a composite map. Because local resolution for NusA-NusG-TEC was superior to that in our previous structure of NusA-NusG-TTC-LC (n = 13; see above), an atomic model was built and refined (procedures as described above).

Cryo-EM data for NusG-TTC-X (n = 12), NusG-TTC-X (n = 13), RfaH-TTC-X (n = 12), and RfaH-TTC-X (n = 13) were collected at the Rutgers University Cryo-EM and Nanoimaging Core Facility, as described above and were processed through the 3D auto-refinement stage as described above (Extended Data Figs. 4-5). For NusG-TTC-X (n = 12), NusG-TTC-X (n = 13), RfaH-TTC-X (n = 12), and RfaH-TTC-X (n = 13), datasets of 3,056, 3,763, 4,586, and 8,012 micrographs, respectively, were recorded from single grids over 2-3 days.

Structure visualization was performed using Coot^51^, Chimera^54^, and PyMOL (Schrödinger).

Final atomic density maps and atomic coordinates for NusA-NusG-TTC-LC (n = 12), NusA-NusG-TTC-LC (n = 13), NusA-NusG-TTC-LC (n = 17), NusA-NusG-TTC-LC (n = 20), NusA-RfaH-TTC-LC (n = 12), NusA-RfaH-TTC-LC (n = 13), NusA-NusG-TTC-B^walked^ (n = 9), and NusA-NusG-TTC-LC^walked^ (n = 13) were deposited in the Protein Data Bank and the Electron Microscopy Data Bank with accession codes PDB 8VOP and EMD-43388, PDB 8VOQ and EMD-43389, PDB 8VOR and EMD-43390, PDB 8VOS and EMD-43391, PDB 8VL1 and EMD-43335, PDB 8VOO and EMD-43387, PDB 9W0N and EMD-65509, and PDB 9Y79 and EMD-72646, EMD-72554, and EMD-72553 (composite map, global map, local map processed by focussed refinement of NusA-NusG-TEC), respectively.

## Extended Data Figure Legends

**Extended Data Fig. 1.**
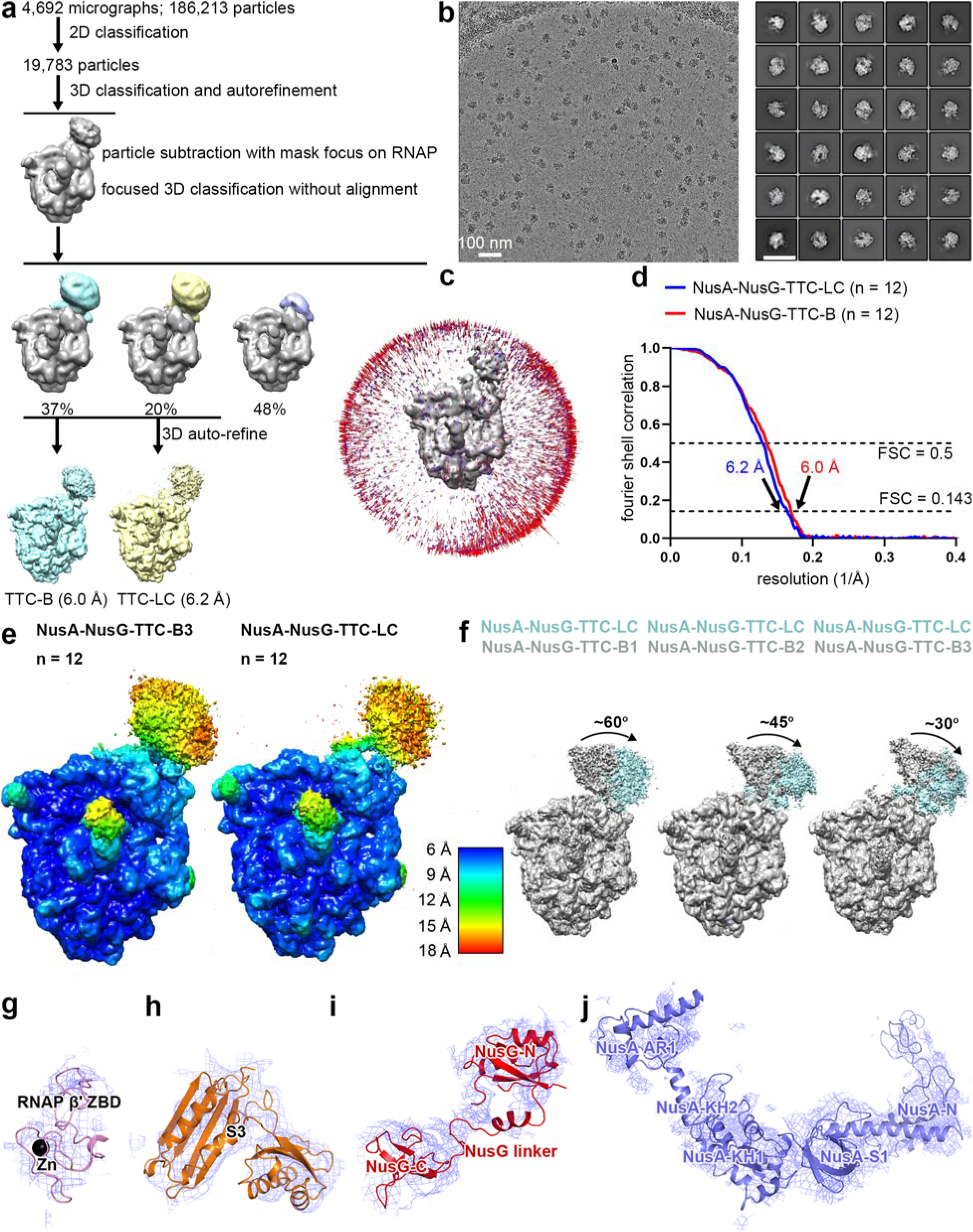
Structure determination: NusA-NusG-TTC-B and NusA-NusG-TTC-LC (n = 12) **(a)** Data processing scheme (Extended Data Table 1). **(c)** Representative electron micrograph and 2D class averages (100 nm scale bar in left subpanel; 50 nm scale bar in right subpanel). **(d)** Orientation distribution. **(e)** Fourier-shell-correlation (FSC) plot. **(f)** EM density maps colored by local resolution. View orientation as in Figs. 1, 2a, and 2b. **(g)** Superimpositions of NusA-NusG-TTC-LC on NusA-NusG-TTC-B subclass B1^32^ (left), subclass B2^32^ (center), and subclass B3^32^ (right). **(g-j)** Representative EM densities (blue mesh) and fits (ribbons) for RNAP β’ ZBD, ribosomal protein S3, NusG, and NusA.

**Extended Data Fig. 2.**
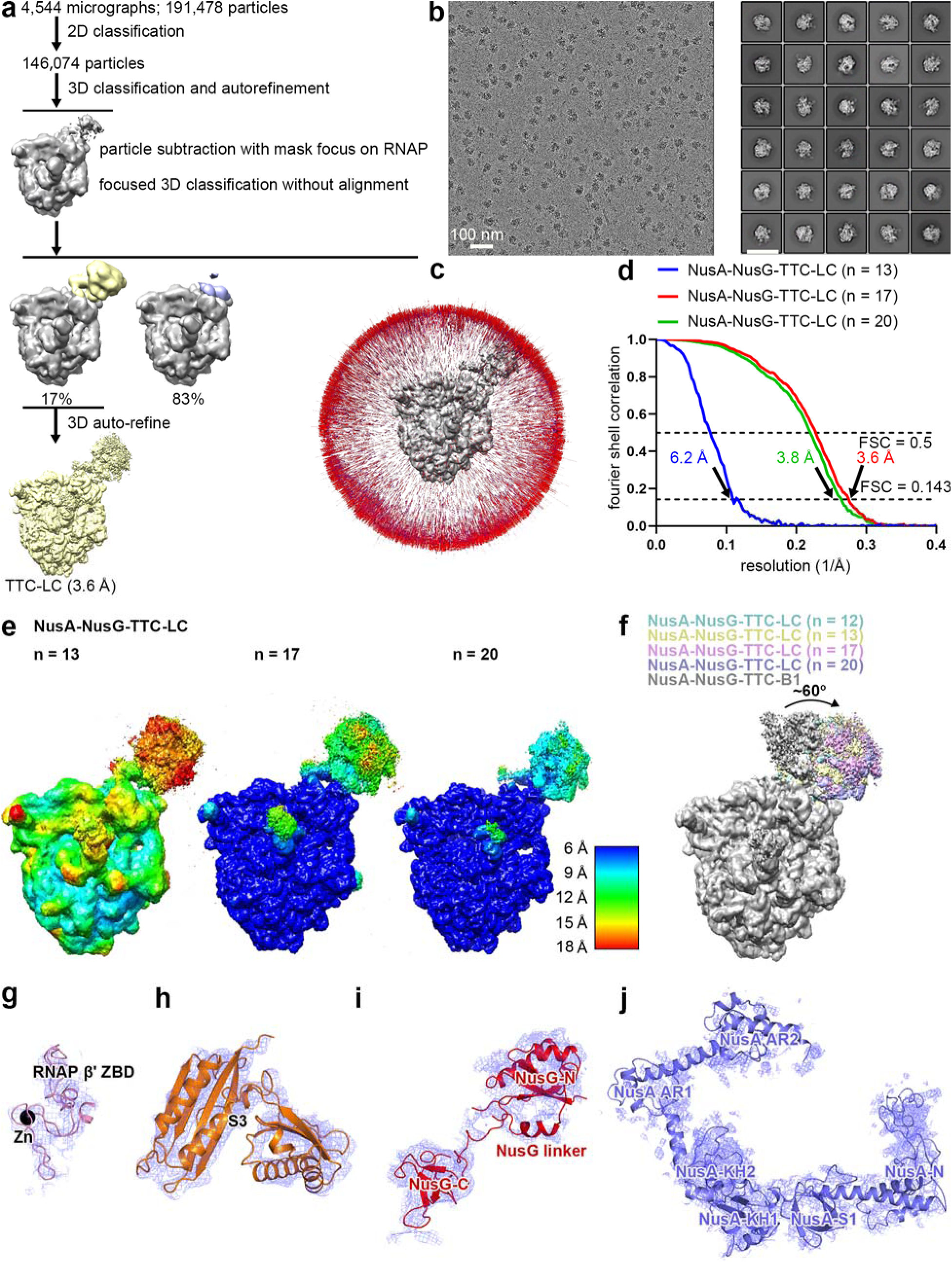
Structure determination: NusA-NusG-TTC-LC (n = 13, 17, or 20) **(a)** Data processing scheme (Extended Data Table 1). **(b)** Representative electron micrograph and 2D class averages (100 nm scale bar in left subpanel; 50 nm scale bar in right subpanel). **(c)** Orientation distribution. **(d)** Fourier-shell-correlation (FSC) plot. **(e)** EM density maps colored by local resolution. View orientation as in Figs. 1, 2a, and 2b. **(g)** Superimpositions of NusA-NusG-TTC-LC (n = 12, 13, 17, or 20) on NusA-NusG-TTC-B subclass B1^32^. **(g-j)** Representative EM densities (blue mesh) and fits (ribbons) for RNAP β’ ZBD, ribosomal protein S3, NusG, and NusA.

**Extended Data Fig. 3.**
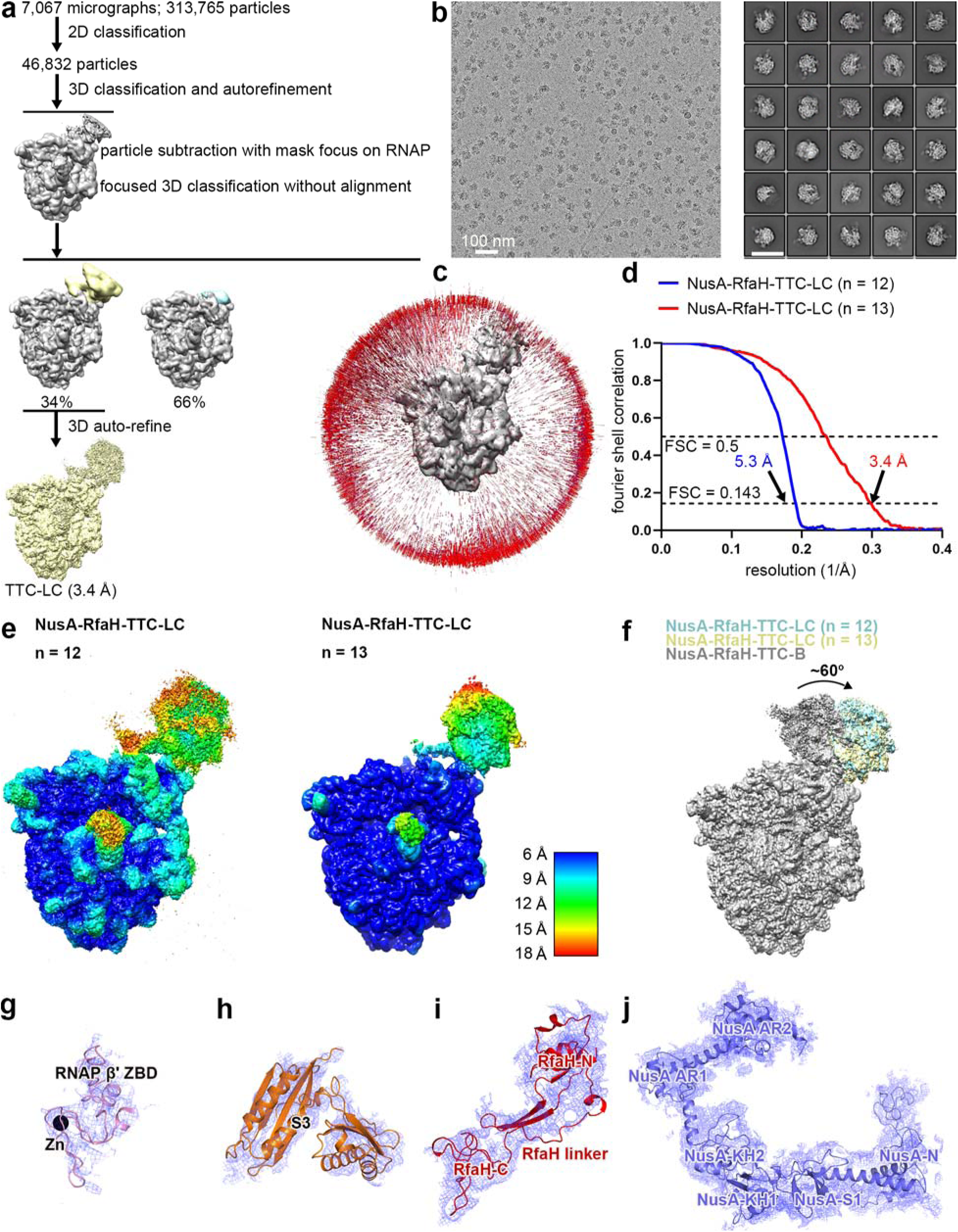
Structure determination: NusA-RfaH-TTC-LC (n = 12 or 13) **(a)** Data processing scheme (Extended Data Table 1). **(b)** Representative electron micrograph and 2D class averages (100 nm scale bar in left subpanel; 50 nm scale bar in right subpanel). **(c)** Orientation distribution. **(d)** Fourier-shell-correlation (FSC) plot. **(e)** EM density maps colored by local resolution. View orientation as in Figs. 1, 2a, and 2b. **(f)** Superimpositions of NusA-RfaH-TTC-LC (n = 12 or 13) on NusA-RfaH-TTC-B^33^. **(g-j)** Representative EM densities (blue mesh) and fits (ribbons) for RNAP β’ ZBD, ribosomal protein S3, RfaH, and NusA.

**Extended Data Fig. 4.**
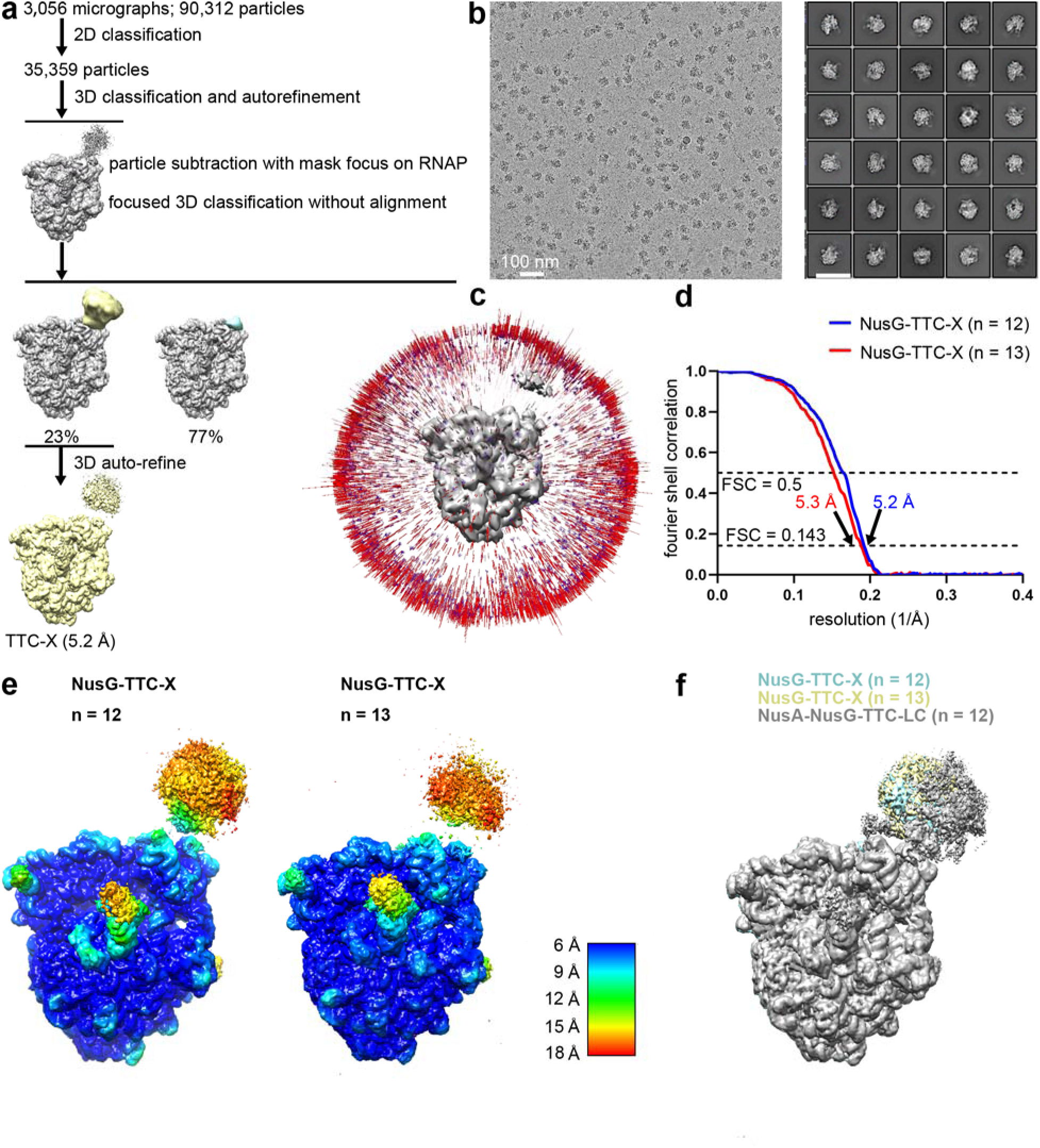
Structure determination: NusG-coupled TTC containing long mRNA spacer in absence of NusA (NusG-TTC-X; n = 12 or 13) **(a)** Data processing scheme. **(b)** Representative electron micrograph and 2D class averages (100 nm scale bar in left subpanel; 50 nm scale bar in right subpanel). **(c)** Orientation distribution. **(d)** Fourier-shell-correlation (FSC) plot. **(e)** EM density maps colored by local resolution. View orientation as in Figs. 1, 2a, and 2b. **(f)** Superimpositions of NusG-TTC-X (n = 12 or 13) on NusA-NusG-TTC-LC (n = 12).

**Extended Data Fig. 5.**
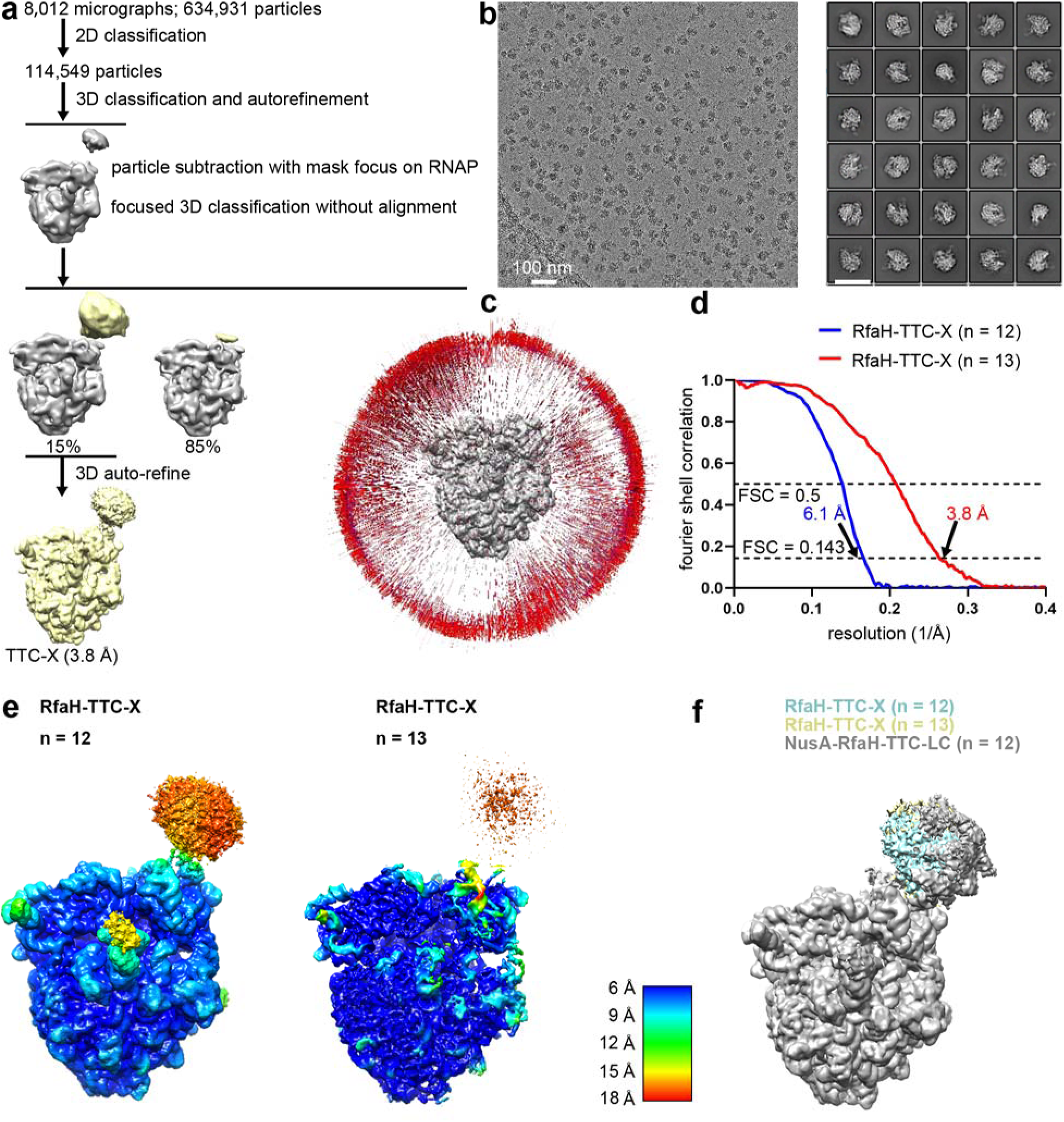
Structure determination: RfaH-coupled TTC containing long mRNA spacer in absence of NusA (RfaH-TTC-X; n = 12 or 13) **(a)** Data processing scheme. **(b)** Representative electron micrograph and 2D class averages (100 nm scale bar in left subpanel; 50 nm scale bar in right subpanel). **(c)** Orientation distribution. **(d)** Fourier-shell-correlation (FSC) plot. **(e)** EM density maps colored by local resolution. View orientation as in Figs. 1, 2a, and 2b. **(f)** Superimpositions of RfaH-TTC-X (n = 12 or 13) on NusA-RfaH-TTC-LC (n = 12).

**Extended Data Fig. 6.**
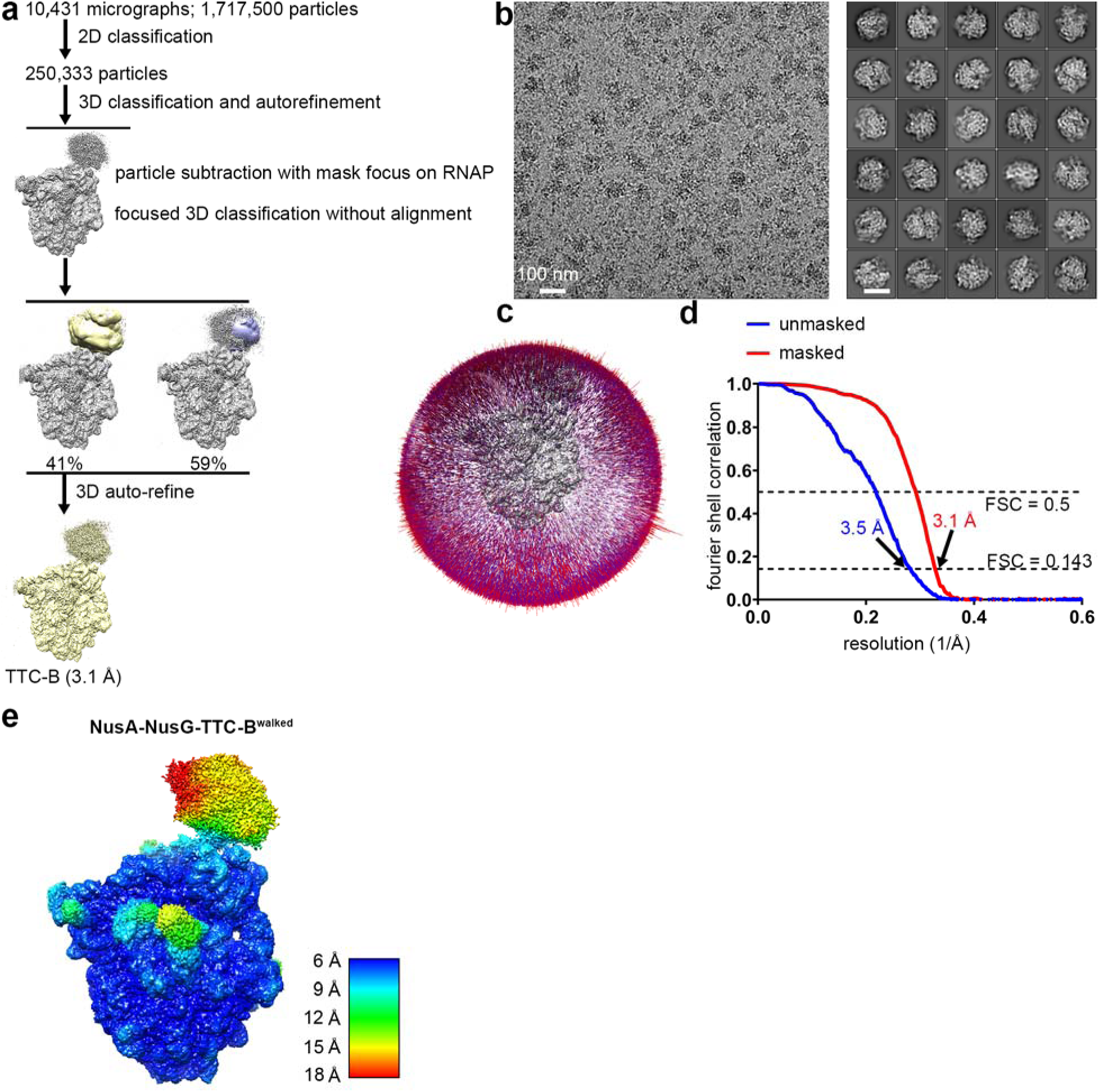
Structure determination: NusA-NusG-TTC-B^walked^ (n = 9; subclass B2) **(a)** Data processing scheme (Extended Data Table 2). **(b)** Representative electron micrograph and 2D class averages (100 nm scale bar in left subpanel; 50 nm scale bar in right subpanel). **(c)** Orientation distribution. **(d)** Fourier-shell-correlation (FSC) plot. **(e)** EM density map colored by local resolution. View orientation as in Figs. 1, 2a, and 2b.

**Extended Data Fig. 7.**
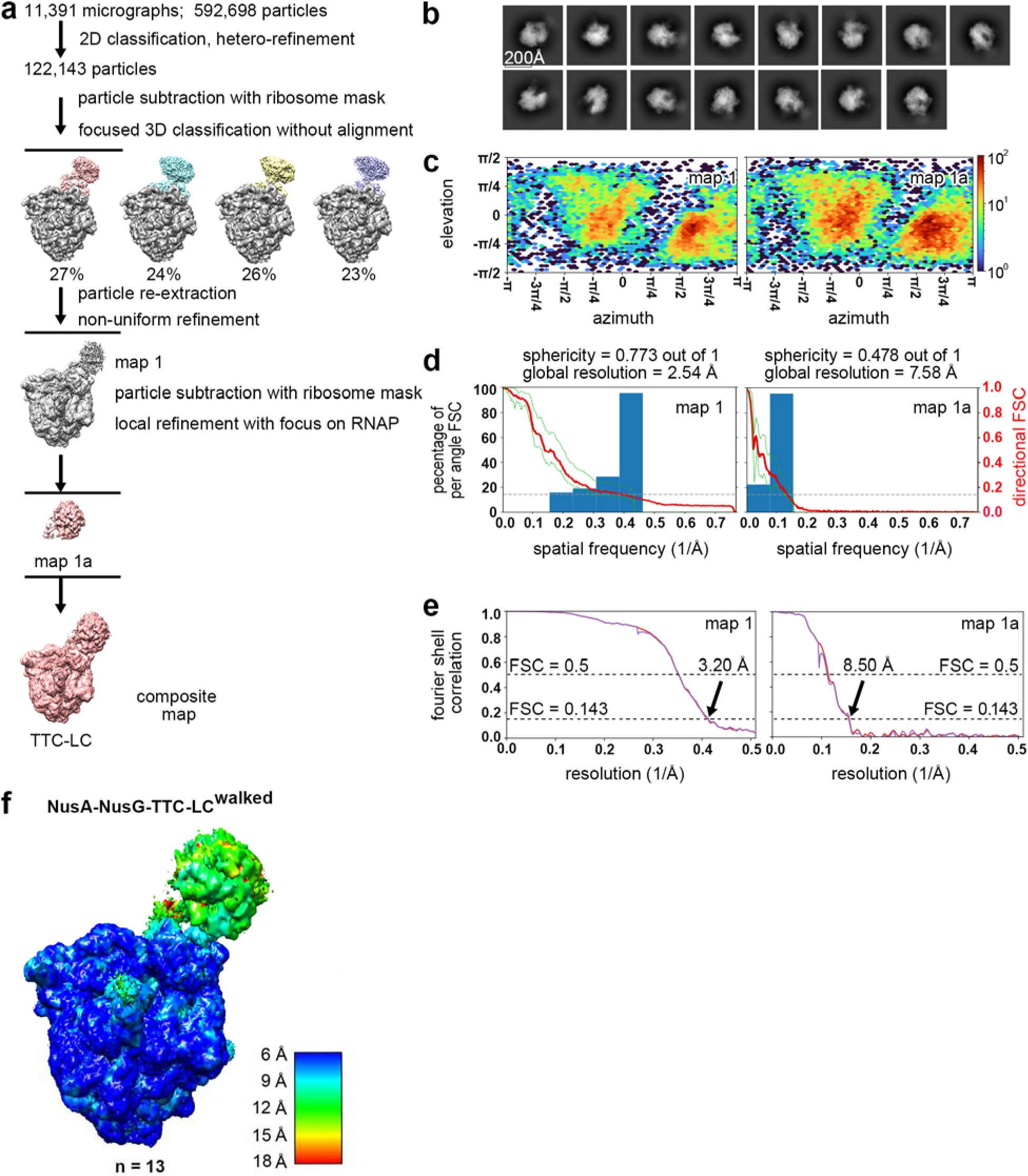
Structure determination: NusA-NusG-TTC-LC^walked^ (n = 13) **(a)** Data processing scheme (Extended Data Table 2). **(b)** Representative 2D class averages. **(c)-(d)** Orientation distribution plots. **(e)** Fourier-shell-correlation (FSC) plots. **(f)** EM density map colored by local resolution. View orientation as in Figs. 1, 2a, and 2b.

**Extended Data Fig. 8.**
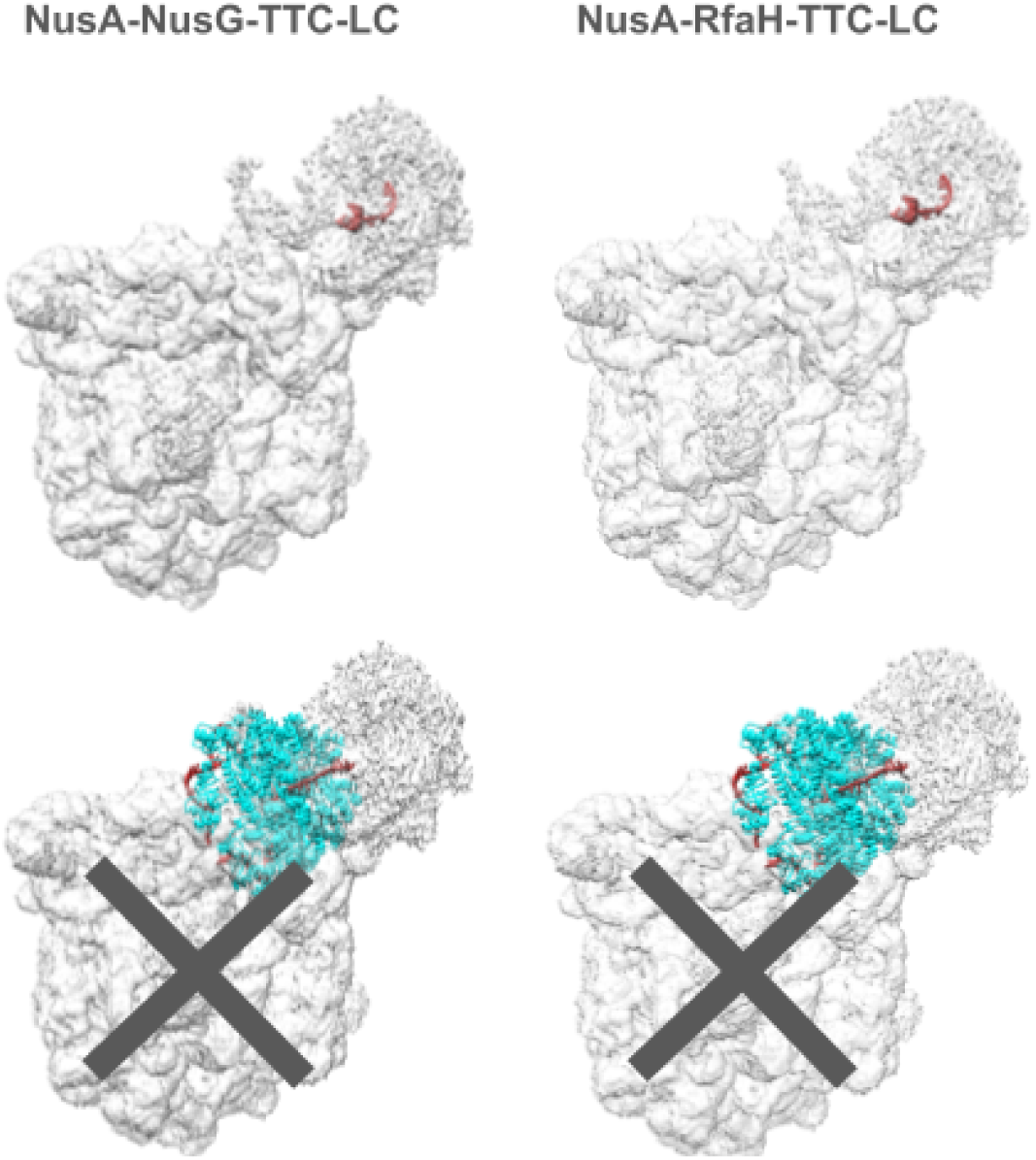
Structural modelling: compatibility of TTC-LC with formation of pause and termination RNA hairpins and incompatibility of TTC-LC with engagement of termination factor Rho. **(a)** TTC-LC (gray) is fully compatible with formation of pause and termination RNA hairpins^40–42^ (red; no steric clash and no interaction of RNA hairpin with the RNA-helicase domain of ribosomal protein S3). **(b)** TTC-LC (gray) is incompatible with engagement of termination factor Rho^24^ (cyan with bound mRNA segments in red; severe steric clash with ribosome).

**Extended Data Table 1:** Cryo-EM structures: NusA-NusG-TTC-B (n = 12) and NusA-NusG-TTC-LC (n = 12, 13, 17, and 20)

|  | NusA-NusG-TTC-B<br>n = 12 | NusA-NusG-TTC-LC<br>n = 12<br>(EMDB-43388)<br>(PDB 8VOP) | NusA-NusG-TTC-LC<br>n = 13<br>(EMDB-43389)<br>(PDB 8VOQ) | NusA-NusG-TTC-LC<br>n = 17<br>(EMDB-43390)<br>(PDB 8VOR) | NusA-NusG-TTC-LC<br>n = 20<br>(EMDB-43391)<br>(PDB 8VOS) |
| --- | --- | --- | --- | --- | --- |
| **Data collection and processing** |  |  |  |  |  |
| Magnification | 130,000x | 130,000x | 130,000x | 130,000x | 130,000x |
| Voltage (kV) | 200 | 200 | 200 | 200 | 200 |
| Electron exposure (e-/Å<sup>2</sup>) | 28 | 28 | 28 | 28 | 28 |
| Defocus range (µm) | -1.25 to -2.0 | -1.25 to -2.0 | -1.25 to -2.0 | -1.25 to -2.0 | -1.25 to -2.0 |
| Pixel size (Å) | 1.038 | 1.038 | 1.038 | 1.038 | 1.038 |
| Symmetry imposed | C1 | C1 | C1 | C1 | C1 |
| Initial particle images (no.) | 1,7581 | 1,7581 | 3,213 | 122,400 | 209,053 |
| Final particle images (no.) | 3,494 | 6,645 | 1,516 | 24,109 | 32,821 |
| Map resolution (Å) | 6.2 | 6.0 | 6.2 | 3.6 | 3.8 |
| FSC threshold | 0.143 | 0.143 | 0.143 | 0.143 | 0.143 |
| Map resolution range (Å) | 5.0-50 | 5.49-50 | 6.13-50 | 3.27-44.37 | 3.29-50 |
| **Refinement** |  |  |  |  |  |
| Initial models used (PDB codes) | 6X6T | 6X6T | 6X6T | 6X6T | 6X6T |
| Model resolution (Å) | 6.3 | 6.6 | 6.8 | 5.8 | 6.0 |

**Extended Data Table 2:**
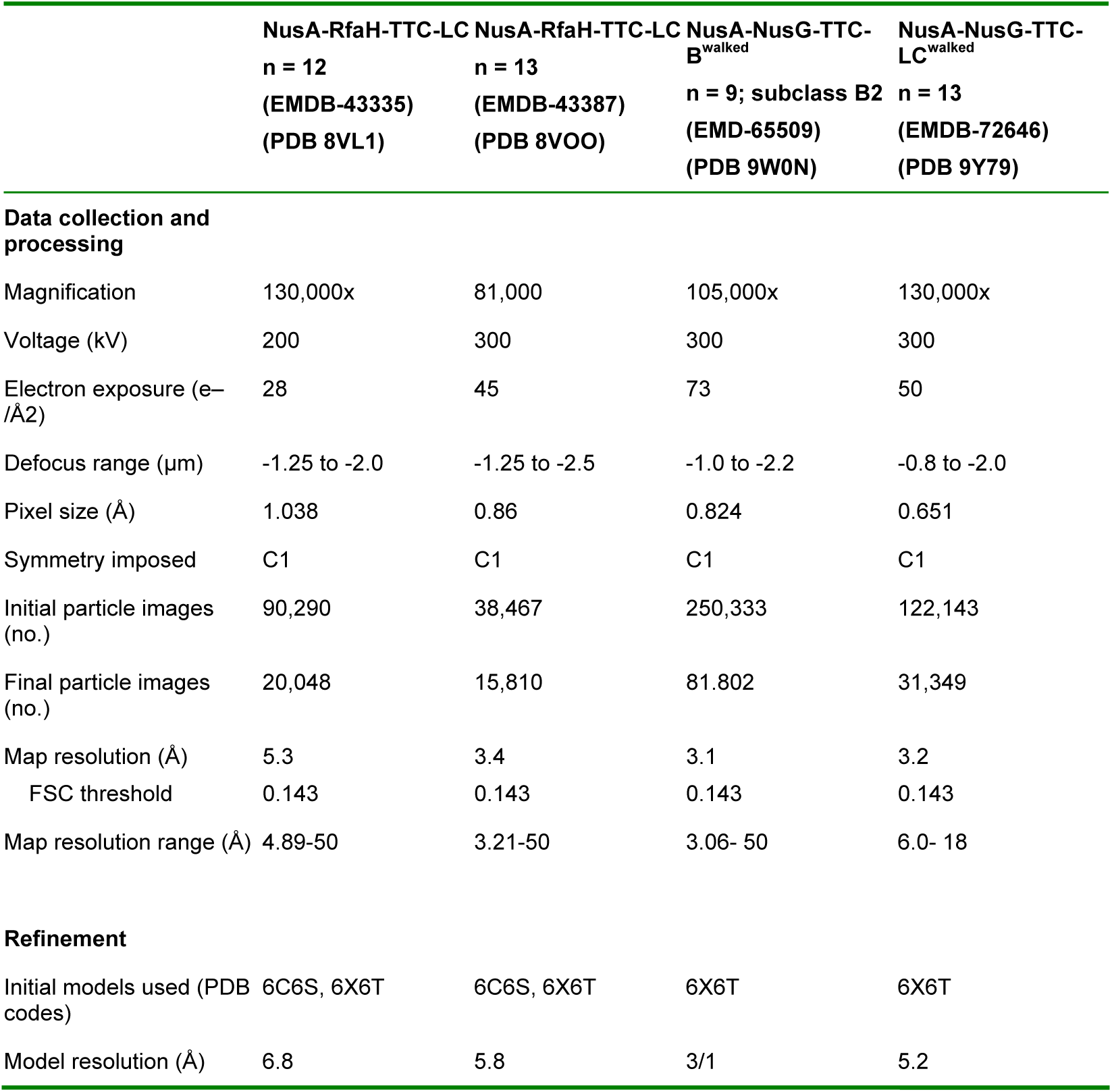
Cryo-EM structures: NusA-RfaH-TTC-LC (n = 12 and 13), NusA-NusG-TTC-B^walked^ (n = 9; subclass B2), and NusA-NusG-TTC-LC^walked^ (n = 13)

